# BIAPSS - BioInformatic Analysis of liquid-liquid Phase-Separating protein Sequences

**DOI:** 10.1101/2021.02.11.430806

**Authors:** Aleksandra E. Badaczewska-Dawid, Davit A. Potoyan

## Abstract

Liquid-liquid phase separation (LLPS) has recently emerged as a cornerstone mechanism underlying the biogenesis of membraneless organelles (MLOs). However, a quantitative molecular grammar of protein sequences that controls the LLPS remains poorly understood. The progress in this field is hampered by the insufficiency of comprehensive databases and associated computational infrastructure for targeting biophysical and statistical analysis of phase separating biopolymers. Therefore, we have created a novel open-source web platform named BIAPSS (BioInformatic Analysis of liquid-liquid Phase-Separating protein Sequences) which contains interactive data analytic tools in combination with a comprehensive repository of bioinformatic data for on-the-fly exploration of sequence-dependent properties of proteins with known LLPS behavior. BIAPSS includes a residue-resolution biophysical analyzer for interrogating individual protein sequences (SingleSEQ tab). The latter allows users to correlate regions prone to phase separation with a large array of physicochemical attributes and various short linear motifs. BIAPSS also includes global statistics derived over the universe of most of the known LLPS-driver protein sequences (MultiSEQ tab) for revealing the regularities and sequence-specific signals driving phase separation. Finally, BIAPSS incorporates an extensive cross-reference section that links all entries to primary LLPS databases and other external resources thereby serving as a central navigation hub for the phase separation community. All of the data used by BIAPSS is freely available for download as well-formatted pre-processed data with detailed descriptions, facilitating rapid implementation in user-defined computational protocols.

TOC - graphical abstract

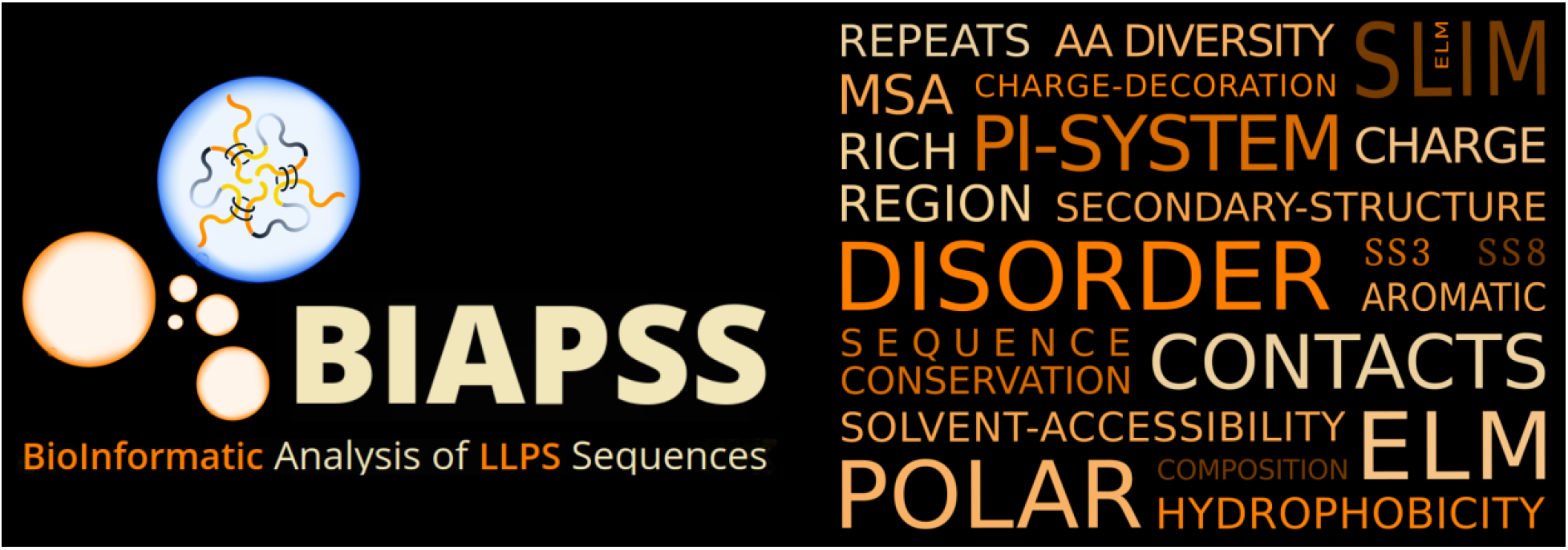

**Author summary:** Proteins, especially those with low complexity and intrinsically disordered regions, have recently come into the limelight because of mounting evidence showing that these regions can drive the formation of membraneless organelles (MLOs) in cells. The underlying physical mechanism for forming MLOs is liquid-liquid phase separation (LLPS); a thermodynamically driven process whereby a cellular milieu with a relatively well-mixed distribution of biomolecules gets decomposed into liquid droplets where the concentration of selected biomolecules is higher. Deciphering molecular sequence grammar of phase separation has turned out to be challenging because of the complexity of this process in cells and the vastness of sequence space of LLPS-driver proteins. While the field is still in its infancy the growth of experimental data has already spurred the creation of several major databases which collect and annotate bimolecular systems with confirmed LLPS behavior. What is currently missing is a framework that would leverage the existing databases by integrating them with deep biophysical and bioinformatic analysis for identifying statistically significant features of protein sequences implicated in LLPS. In this work, we have addressed this challenge by creating an open-source web platform named BIAPSS (BioInformatic Analysis of liquid-liquid Phase-Separating protein Sequences) which integrates a comprehensive repository of pre-processed bioinformatic data for LLPS-driver protein sequences with interactive analytic applications for on-the-fly analysis of biophysical features relevant for LLPS behavior. BIAPSS empowers users with novel and effective tools for exploring LLPS-related sequence signals for individual proteins (SingleSEQ tab) and globally by integrating common regularities across subgroups or the entire LLPS sequence superset (MultiSEQ). The long-term plan for BIAPSS is to serve as a unifying hub for the experimental and computational community with a comprehensive set of analytic tools, biophysically featured data, and standardized protocols facilitating the identification of sequence hot spots driving the LLPS, which all can support applications for designing new sequences of biomedical interest.

## Introduction

In the past few years, the liquid-liquid phase separation (LLPS) of biomolecules has become a universal language for interpreting intracellular signaling, compartmentalization, and regulation[1–5]. The ability to phase separate appears to be encoded primarily in the protein sequences, frequently containing disordered and low complexity domains, which are enriched in charged and multivalent interaction centers[6–8]. Nevertheless, the quantitative aspects of how amino acids encode and decode the phase separation remain largely unknown[9–11]. This is because many different combinations of relevant interactions seem to be contributing to phase separation without anyone being universally necessary[12]. So far, however, with a few exceptions[13–16] mostly case-by-case studies of different sequences are performed, with the broader context of many findings, including their statistical significance remaining unknown.

In recent years there appeared many databases that collect LLPS-related biomolecular complexes. Some of the prominent examples are the PhaSepDB[14], PhaSePro[15], LLPSDB[16], and DrLLPS[17]. These databases enable browsing and downloading of entries and provide detailed information on experimental conditions for bimolecular condensation. Above all, these databases collect and annotate partially overlapping sets of phase-separating protein sequences. In-particular PhaSePro, LLPSDB, and a subset of PhaSepDB contain manually curated protein sequences recognized to directly drive the formation of subcellular compartments. Accumulation of high-quality datasets is certainly a necessary condition for making progress towards uncovering driving forces of protein phase separation. Specifically, these carefully curated sequences can be used in the comprehensive bioinformatic inquiry to identify and analyze a set of biophysically motivated molecular features of protein phase separation. Nevertheless, there is still a need for an overall perspective integrating single features and observations into a merged characteristic.

Further improvement can be made by combining all of the known biosystems from the primary LLPS databases and providing easy tracking of corresponding entries in the external sources, such as UniProt (protein sequences)[18], DisProt (disordered proteins)[19], Protein Data Bank (protein structure)[20], Compartments (cellular location)[21], and others. Furthermore, the comprehensive analysis of biophysical properties and detection of various short linear motifs can speed up the discovery of statistically significant sequence signals for proteins with LLPS behavior. To this end, we have developed a web platform BIAPSS: BioInformatic Analysis of liquid-liquid Phase-Separating protein Sequences, available online at https://biapss.chem.iastate.edu/. The main objective of BIAPSS is to enable a rapid and on-the-fly deep statistical analysis of LLPS-driver proteins using the pool of sequences with empirically confirmed phase behavior.

## Overview of BIAPSS web platform

### BIAPSS content and data sources

BIAPSS is designed as a user-friendly web platform that is billing itself as a central resource for systematic and standardized statistical analysis of biophysical characteristics of known LLPS sequences. The web service provides users with (i) a database of the superset of experimentally evidenced LLPS-driver protein sequences, (ii) a repository of pre-computed bioinformatics and statistics data, and (iii) two sets of web applications supporting the interactive analysis and visualization of physicochemical and biomolecular characteristics of LLPS proteins. The initial LLPS sequence set leverages the data from manually curated primary LLPS databases, namely PhaSePro[15] and LLPSDB[16]. Given that the number of experimentally confirmed LLPS driver proteins is constantly growing, the BIAPSS pre-computed repository will be updated annually and released to the public. This will significantly save the users time by eliminating the need for exhaustive in-house calculations. The newly developed interactive web applications integrate the results from our extensive studies, described in more detail elsewhere [63].

### BIAPSS web services and features

The layout and main functionalities of BIAPSS services are summarized in Figure 1. The general outline of the platform is designed to provide clarity and intuitive navigation by avoiding the excess of permanently visible information. Due to the multitude of analyses, available to meet the needs of a diverse audience of scientists, the extensive content of BIAPSS has been divided into 5 main tabs.

**Figure 1.**
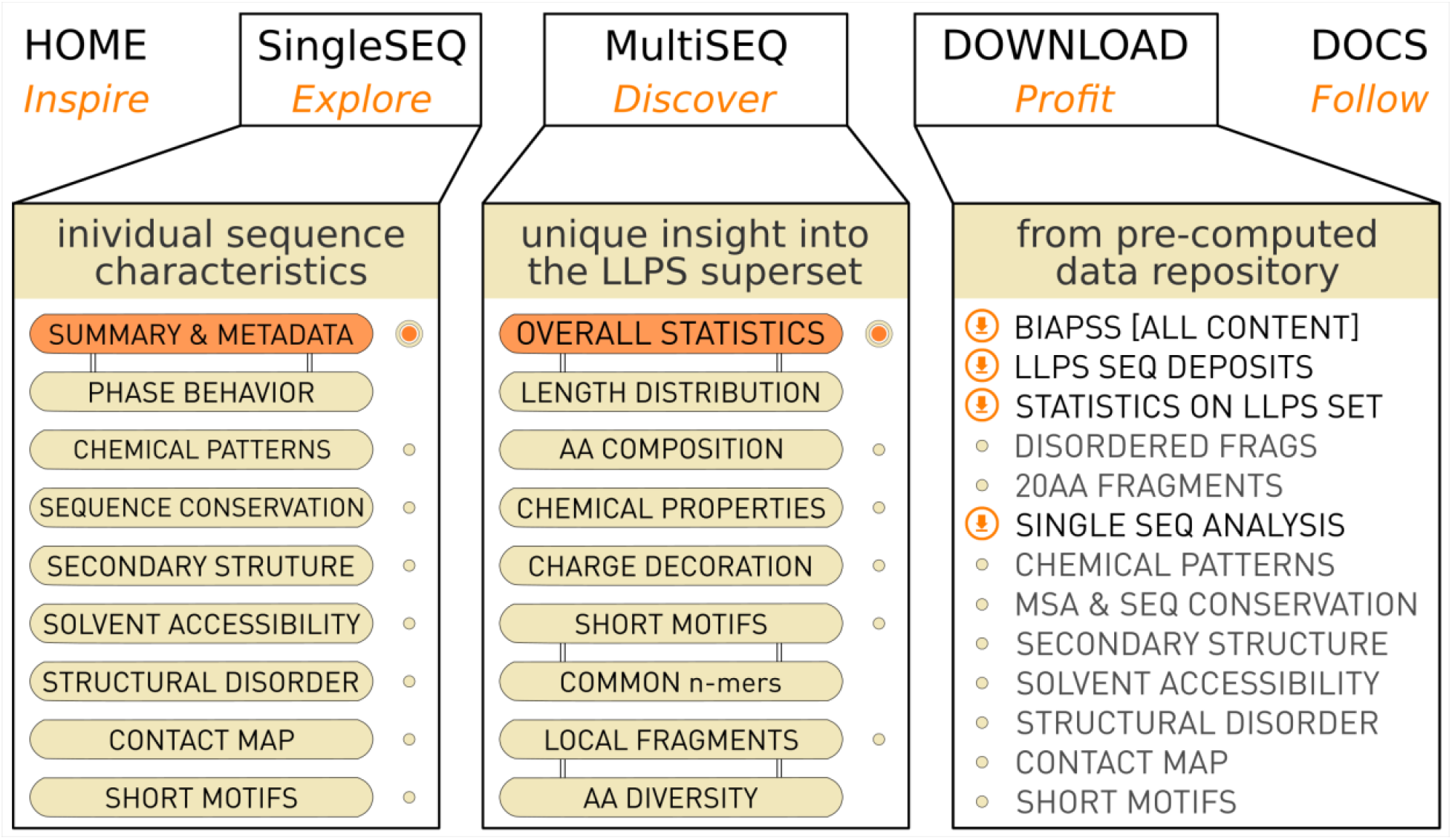
The overall layout of BIAPSS web platform (https://biapss.chem.iastate.edu) for comprehensive sequence-based analysis of LLPS proteins. The core of the implemented web applications and data repository is contained in the SingleSEQ, MultiSEQ, and Download tabs.

The Home tab is a place where the user gets a high-level overview of the features of BIAPSS services. Next comes the SingleSEQ tab which is dedicated to the exploration of individual LLPS sequence characteristics. Besides a case summary and cross-reference section, there are multiple web applications dedicated to the in-depth analysis of biomolecular features, such as sequence conservation with multiple sequence alignment (MSA)[22], various sequence-based predictions by the state-of-the-art methods for secondary structure[23–27], solvent accessibility[26–28], structural disorder[26,29–33], contact maps[26,31,33], and uniquely proposed detection of numerous short linear motifs (SliMs)[34–38] recently highlighted as key regions for driving the LLPS[39]. The MultiSEQ tab provides the user with a set of web applications for a broad array of statistics on a superset of LLPS sequences. One may there investigate the regularities and trends specific only for disordered regions, such as amino acid (AA) composition, including AA diversity or regions rich in a given AA, general physicochemical patterns of polarity, hydrophobicity, the distribution of aromatic or charged residues, including not only the overall net charge but also charge decoration parameters that emerged as a relevant factor for electrostatic interactions of intrinsically disordered proteins (IDPs)[40], and more. Also, a deeper focus on the general frequency of particular short linear motifs, including LARKS[35], GARs[36], ELMs[34], and steric zippers[38], as well as pioneering identification of specific n-mers, can bring new perspectives in the field. The Download tab facilitates accessing the BIAPSS repository. The available data includes raw predictions pre-calculated using the well-established tools as well as the findings of our deep statistical analysis. For the convenience of users, we have unified and integrated the pre-processed results into a standardized CSV format accompanied with intuitive descriptors to facilitate reuse and, specifically, allows the researcher to implement the pre-computed data directly or carry out further analysis. Finally, in the Docs tab, the user can follow the detailed data-analytic workflow and learn more about used tools with corresponding references to the original literature. The documentation also includes an easy-to-use tutorial dedicated to individual web applications, where all of the features are presented graphically with detailed descriptions (see also Supplementary Information, where the user manual is attached).

## Design and implementation

### Full-stack development in Python with Plotly-Dash graphing library

The back-end processing pipeline of BIAPSS is implemented in the Python framework, where in-house developed algorithms parse pre-computed data and perform on-the-fly analysis. The basic front-end user interface of the BIAPSS web platform is implemented with the HTML5, CSS, JavaScript, and Bootstrap components which support the responsiveness and mobile-accessibility of the website. Specifically, our cross-platform framework is adjusted to be run on multiple operating systems and popular browsers. Modern display-layer solutions improve user experience by enabling smooth loading of contents, page transitions, and accompanying an in-depth presentation of the results. For instance, we included a lightbox slideshow with a brief overview of the features, collapsed menu, and modal images of quick guide within individual applications, side navigation, and more. Interactive graph plotting and data visualization accessible through web applications in SingleSEQ and MultiSEQ tabs were developed with the Plotly-Dash[41] browser-based graphing libraries for Python which create a user-responsive environment and follow remote, customized instructions. Thanks to the interactive interface users can go directly from exploratory analytics to the creation of publication-ready high-quality images.

### Organization and visualization within BIAPSS web applications

The characteristics of an individual protein sequence can be explored on-the-fly either through web applications or by downloading and doing further processing according to the user's needs. Each category of analysis matches a separate application. The switching between applications is possible by both scrolling-down and through a user-friendly navigation panel (expandable on the center-right), which automatically centers the page on the selected app. To make it intuitive and easy-to-use, the overall layout is unified for all the applications and shown in Figure 2.

**Figure 2.**
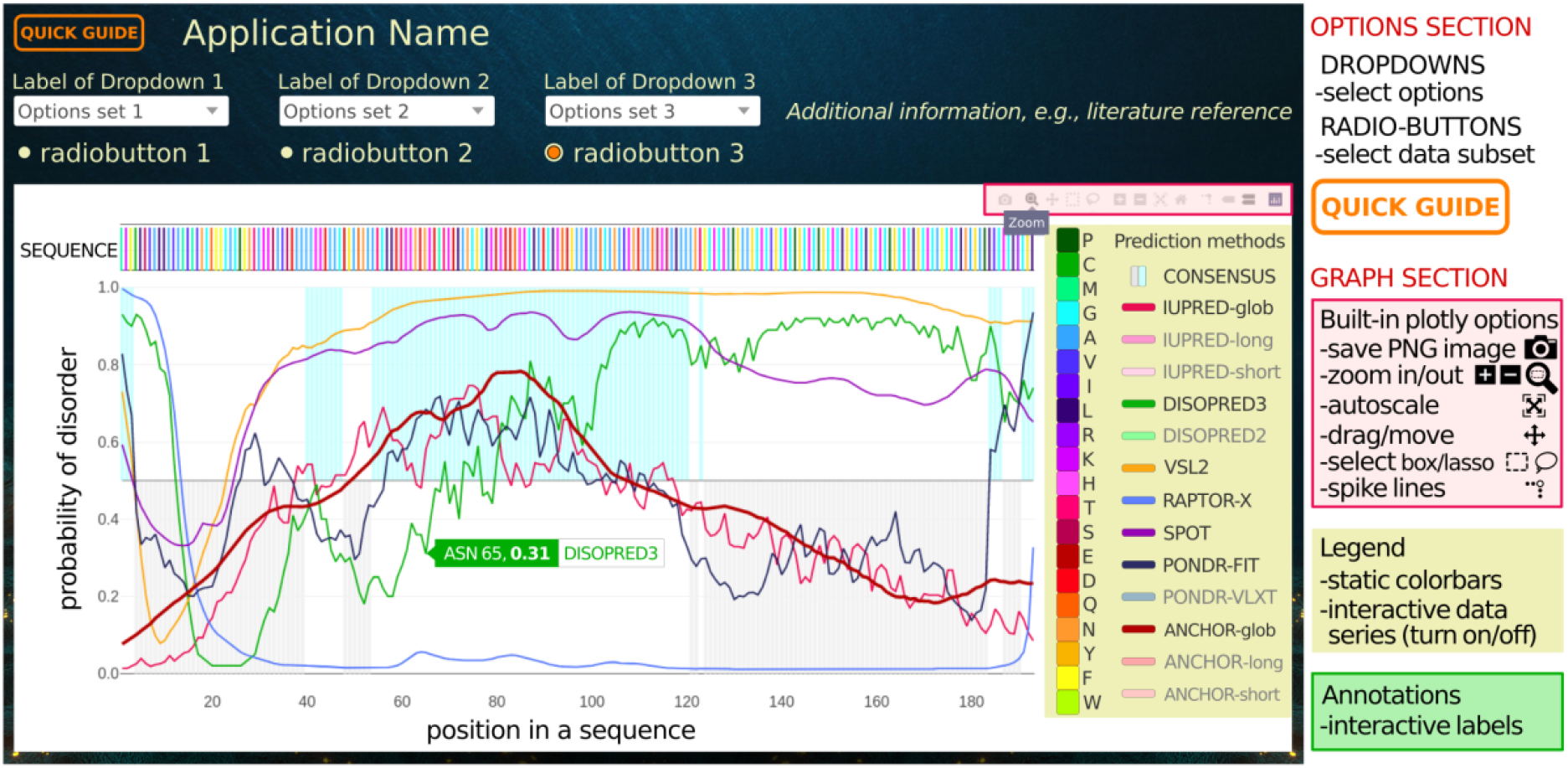
The unified layout of developed interactive web applications available in *SingleSEQ* and *MultiSEQ* tabs of the BIABSS platform. The application name specifies the type of analysis. The *Quick Guide* button in the upper left corner displays a simplified tutorial. The app contains option and graph sections. Built-in Plotly options allow for interactive graph modification and saving.

Each individual web application is running in the single-page framework which is automatically adjusted to the user viewport, including mobile devices with minimal dimensions being 300×600 px. The app’s name, indicating the general type of analysis, is provided on top of the page. On the top-left, an orange “quick guide” button displays a brief tutorial. The options section typically includes dropdowns and radio-buttons with self-explaining labels or tooltips, that facilitate the customization of the analysis. Querying coupled with dynamic filtering allows the user to explore the LLPS sequences by taking custom slices of the data. This is done by graphing the results from a chosen tool, selecting the parameter along which the data is projected, switching between data subsets, and indicating the reference dataset. The multi-level option section provides a more flexible exploratory of the data following the user’s interests. The graph's layout typically contains multiple rows, where different characteristics are shown along the amino acid sequence. This makes it easier to spot valuable trends even for non-expert users. Each position in the sequence is labeled interactively with detailed analysis-specific annotations. The legend is usually provided on the right, including (i) static color scales assigned to various parameters (ii) and an interactive list of data series that determines which traces are visible on the graph. On the top right corner, the built-in plotly options offer several tools for interactive activity with a graph, including zoom in/out, autoscale, drag/move, select fragment, spike lines, and more. Also, the graph or tailored slice can be downloaded as high-resolution PNG images, ready for publication.

## Results

### Analysis of an individual LLPS sequence

Our deep bioinformatical analysis of individual LLPS protein sequences led to the featurization of biomolecular characteristics into eight categories which are outlined in Figure 3. Each category matches a separate web application implemented within the **SingleSEQ** tab of the BIAPSS platform, https://biapss.chem.iastate.edu/single_seq.html. In the following paragraphs, we list all of the apps by pointing out their aims and available functionalities. More detailed descriptions can be found in the user manual provided in the Supporting Information file.

**Figure 3.**
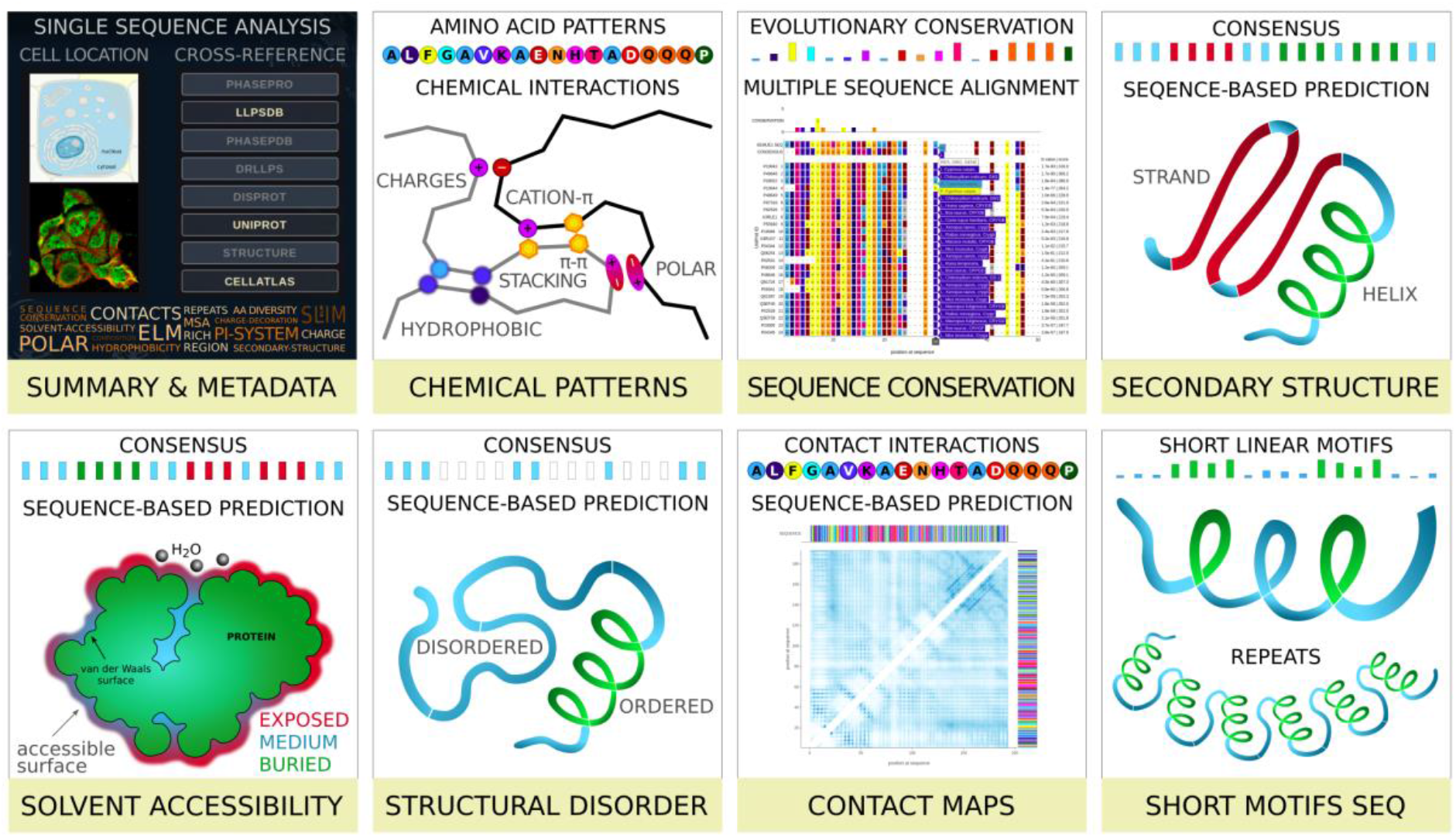
Graphical overview of the types of analyses employed in BIAPSS SingleSEQ web applications for studying individual protein sequences with confirmed LLPS behavior.

### Summary and Metadata web application

First, we provide a compact one-page summary for a single LLPS protein sequence accompanied by rich metadata. A brief overview of all key parameters helps to get an insight into the essence of individual sequence characteristics. The extended Cross-References section integrates many external resources (without embedding their content) and emerges as a central navigation hub for the phase separation-related scientific community. Over there, the user can (i) easily follow the corresponding entries in the primary LLPS-related databases, such as PhaSepDB[14], DrLLPS[17], LLPSDB[16], and PhaSePro[15], (ii) explore the principal annotation data and functional information in UniProt resource[18], (iii) identify manually curated intrinsically disordered regions in DisProt database[19], (iv) check subcellular location through Compartments[21] and see (v) experimental mapping in The Human Protein Atlas[42]. If an experimental structure is known, (vi) the Structure button links to the PDBe-KB repository[43], otherwise the computationally modeled structure is accessible via the MODBASE database[44] and/or SWISS-MODEL server[45]. Finally, the central section contains a summary of individual LLPS sequences, including PSPredictor score estimating the propensity to phase behavior[46], empirically determined LLPS-related region(s), and PubMed identifiers referring to the literature.

### Patterns of Chemical Decoration web application

The Chemical Patterns application allows combining several key biophysical properties for an individual LLPS-driver protein sequence, all within a single multi-row chart. The selected sequence is color-coded according to the individual amino acids and is ordered by their chemical nature. The next 4 rows give an insight into the evolutionary conservation of the protein sequence, providing the consensus sequence profile of multiple sequence alignment and additional MSA-based parameters such as attribute, diversity, and strength. All of this helps to identify the regions consolidated by evolution that are indicative of functional or structural similarity, including the location of insertions and deletions. In the next few rows, there are patterns of chemical properties such as polarity, hydrophobicity, aromaticity, and charge. Then, the robust consensus predictions of secondary structure, solvent accessibility, and structural disorder are provided. The position-based vertical alignment of all features gives a deeper glimpse into the correlation of chemical decoration and structural properties. Furthermore, the all-in-one chart facilitates the detection of sensitive regions for key interactions. For example, parallel comparison of specific patterns of charged or aromatic residues and unique insertion in a structurally disordered region helps to identify hot spot residues or short motifs which may play an important role in the LLPS behavior.

### Sequence Conservation web application

The conservation strength of a sequence region may be a consequence of its functional or structural importance. Therefore, the unique insertions, deletions, or substitutions, identified within the LLPS protein sequence may prove to influence the chemical properties of the polypeptide chain towards multivalent interactions. The Sequence Conservation application delivers (i) the multiple sequence alignment of the LLPS sequence against the reference databases of protein sequences, (ii) the consensus sequence profile, (iii) and evolutionary conservation at the level of individual amino acids. The MSA section contains HMMER-aligned[22] protein sequences colored by the chemical character of amino acids and sorted by the E-value (score is also provided). Each position in the sequence is annotated interactively by an index at the alignment profile, amino acid, organism, and gene.

### The Secondary Structure web application

Conformational preferences of local sequence fragments are conditioned by the local composition of amino acids through the formation of regular patterns of hydrogen bonds along the protein backbone. Recent findings imply that some intrinsically disordered fragments of the sequence tend to get structurally organized into steric zippers or aromatic-rich kinked β sheets [35,38,47,48]. The most advanced sequence-based secondary structure predictors provide not only the binary assignment of ordered regions but also the probability fractions for the helical (H), extended (E), or coiled (C), given for an individual amino acid along the sequence[49,50]. This opens up the possibility to determine factors supporting multivalent interactions as well as detecting regions slightly prone to form ordered structure which can be acting as a switch depending on the environment conditions and molecular crowding. The Secondary Structure application presents the sequence-based predictions of secondary structure elements assigned along with the selected LLPS sequences. Our benchmarking approach combines 5 well-established methods, i.e., PSIPRED[27], RaptorX[26], PORTER-5[26], SPIDER-3[24], FESS[25], and aligns their results per position in the sequence within a multi-row graph. We also provide an agreed consensus of all methods that makes the final result more reliable. In the last section, the detailed probabilities to form each type of HEC element are attributed along the sequence with amino acid resolution.

### The Solvent Accessibility web application

The Solvent Accessibility application presents the sequence-based prediction of structurally buried or exposed regions along the protein sequence. Since the structural properties of many phase-separating proteins are hardly graspable, the analysis of surface solvent accessibility may support, to a certain extent, the identification of regulatory regions prone to temporarily stabilized interactions at the contact interface[51,52]. Our comparative approach matches the results of three state-of-the-art predictive tools, namely RaptorX[53], PaleAle5[27], and SPOT-1D[28]. For compatibility and user-friendliness, the overall layout of the interactive graph is similar to the Secondary Structure app and contains three main sections: (i) on the top, the agreed consensus of solvent accessibility predictions provided in residue-resolution, (ii) followed in the next few rows by the original results derived and unified from the individual methods, (ii) below which there is a section of detailed probabilities for an expose or burial of individual residues along the sequence. The interactive annotations include residue index, amino acid type, and solvent accessibility in three-state notation or particular probabilities.

### The Structural Disorder web application

Identification of low-complexity and unstructured fragments within proteins has recently gained prominence, following the discovery that the liquid-liquid phase separation emerges as a common functioning mechanism for partially or fully disordered proteins[1,2,4,5]. From these findings, we know that protein sequences not only encode for well-ordered tertiary structures but can also explicitly encode intrinsic disorder. The Structural Disorder application provides predictions of disordered regions resulting from the benchmark of numerous widely used methods, including IUPred2A[33] (glob, short, long), DISOPRED2[54], DISOPRED3[32], VSL2[30], RaptorX[53], SPOT-Disorder[31], PONDR-FIT[29], PONDR-VLXT[55]. Most of these methods return the probability of disorder for each position in the sequence. Usually, the residue is considered as ordered when the score is below 0.5. Within the app, the probability section displays the probability distributions of disorder along the sequence for all methods selected by the user. The agreed outcome of 8 well-established tools results in a simplified binary consensus of structural disorder assigned per amino acid position. Accordingly, the ANCHOR[56] method was used for predicting protein binding regions in disordered fragments.

### Contact Map web application

The Contact Map application provides sequence-based predictions of spatial contacts between residues distant in a protein sequence. The most powerful contact predictors, such as those selected for this work are RaptorX Contact[57], SPOT-Contact[28], and ResPRE[33] methods which employ the convolutional neural networks, machine learning, and evolutionary coupling techniques to prepare the list of residue pairs occurring close enough in space with a certain probability. An in-depth analysis of the contacts predicted for the LLPS proteins, despite their largely disordered nature, can help identify regions essential for maintaining structure, dynamics, function and may indicate a lead relevant to phase behavior. The app allows for plotting the square heatmap, colored according to contact probabilities of the selected method. Usually, a score above 0.5 is deemed to be a reliable contact. The interactive annotation shows the detailed probability of contact between the pair of interacting residues.

### Short Linear Motifs web application

The Short Motifs SEQ application is designed to detect various short linear motifs in the individual LLPS sequences. The findings are displayed on the background of regions experimentally confirmed as LLPS-related and/or were predicted to be structurally disordered. The evolutionary conservation of SliMs within intrinsically disordered fragments clearly points to their significance for the regulatory function, hence may also be essential for the collective formation of MLOs. The multi-row application offers a broad screening of short motifs known from the literature as relevant for phase behavior, such as LARKS[35], glycine-arginine-rich regions[36], steric zippers[38], and S--D/S--E motifs specific for phosphosites[37]. Accordingly, the ELMs[37], experimentally validated in eukaryotic cells, are indicated along the sequence with a discrimination of the main ELM classes. Additionally, short fragments frequently repeated along an individual sequence are detectable, including random and tandem repeats. The detailed information on the location, motif class, and instance are displayed on the interactive label.

### Statistics on LLPS sequence superset

Our comprehensive statistics performed on the LLPS-driver sequences superset provides a broad overview of general biophysical characteristics and regularities common for known phase separating protein sequences. This makes it possible to infer certain overall regularities that may be helpful not only to identify efficiently phase-separation-prone sequences but also in designing modifications that favor biomedical applications. The available analyses are outlined in Figure 4 and can be explored within the **MultiSEQ** tab of the BIAPSS platform, https://biapss.chem.iastate.edu/statistics.html. In the following paragraphs, we list all of the statistical analyses by pointing out their aims and available functionalities. The user manual provides graphical snapshots and detailed descriptions which can also be found in the Supporting Information file.

**Figure 4.**
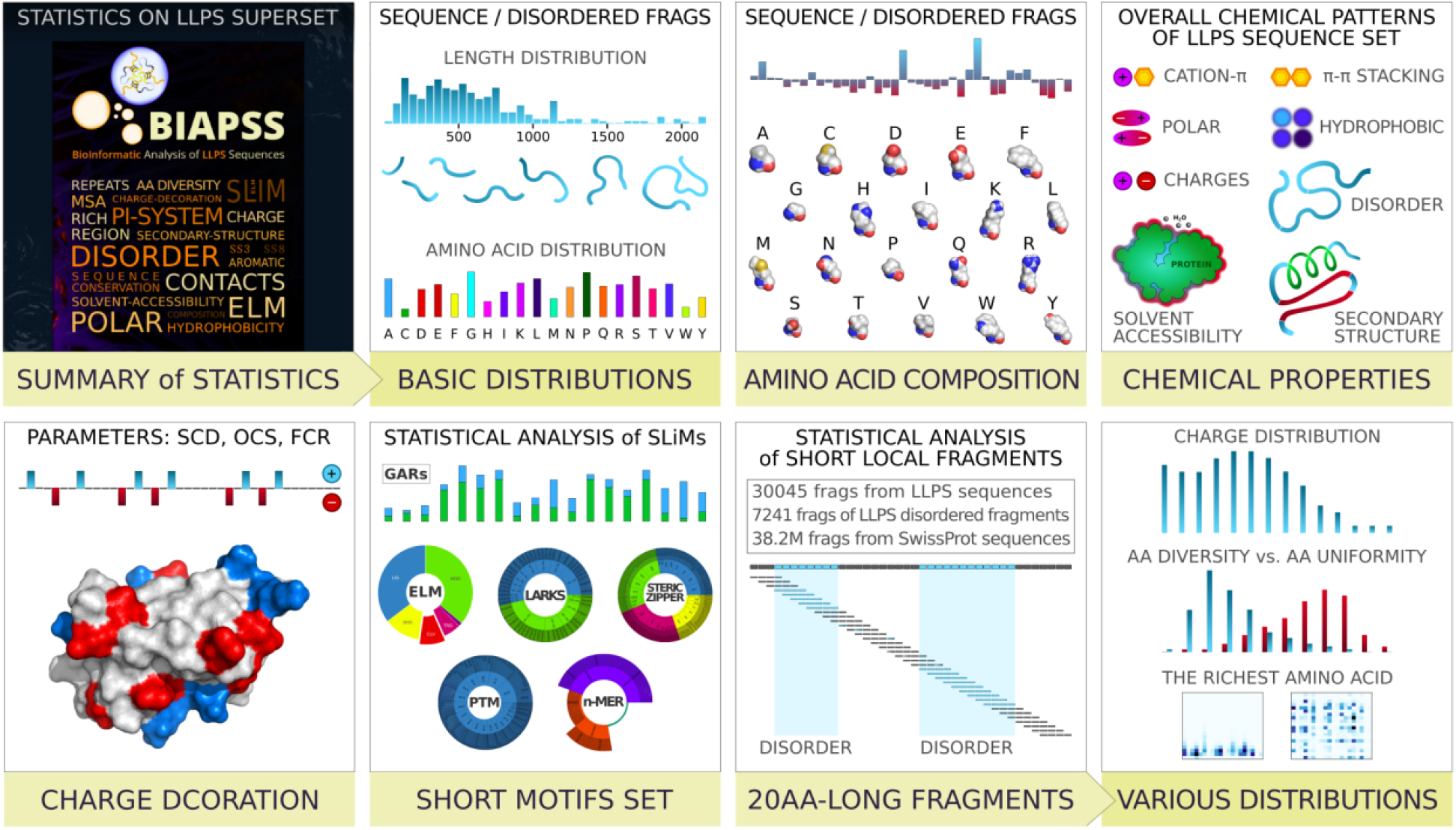
Types of analyses designed to study statistically significant regularities observed across LLPS protein sequences superset.

### Overall Summary of Statistics

The first provided section serves as the brief summary of various statistical analyses performed on the complete set of experimentally confirmed LLPS-driving proteins. A broad overview includes general biophysical properties such as polarity, hydrophobicity, aromaticity, charge split into a positive and negative fraction. The overview also includes some structural characteristics, such as secondary structure (split into helical and extended), solvent accessibility (split into buried and exposed), and structural disorder. The recently proposed charge decoration parameters, i.e., sequence charge decoration (SCD)[58] and overall charge symmetry (OCS)[59], together with a fraction of charged residues (FCR), are provided. Also, there is an actual number of BIAPSS-deposited LLPS-driver protein sequences, the content of disordered regions, the number of organisms of origin, and a subset of human proteins. Such a global overview of biophysical characteristics gives a quick high-level glimpse into the LLPS protein set compared to other reference sets. The deeper analysis is provided in the following more specific interactive applications. In separate tabs within the same category, the user can find the *Length Distribution* and *Amino Acid Distribution* and simultaneously compare the statistics performed for the complete LLPS sequences and extracted disordered regions.

### Amino Acid Composition Analysis

The Amino Acid Composition application enables the analysis of amino acid composition on the superset of LLPS-driver protein sequences. In the interactive graph, the user can compare fractions of all 20 biogenic amino acids simultaneously for all LLPS-driver protein sequences deposited in the BIAPSS repository. The attractive heatmap type of plotting allows for a compact and straightforward all-in-one view which highlights the protein sequences exceedingly rich in a given type of amino acid relative to the complete LLPS set. Additional customization facilitates comparison with the benchmarked statistics derived for other protein groups segregated by the organism or protein type (globular, transmembrane, disordered)[60,61]. This supports the identification of trends in amino acid composition specific for phase-separating or disordered proteins.

### Chemical Properties Analysis

The Chemical Properties application collectively summarizes the chemical properties and biomolecular characteristics on a single chart, for all the available LLPS sequences in the BIAPSS repository. Specifically, the user can investigate the overall distribution of various chemical properties, such as polarity, hydrophobicity, charge (total, positive, negative), aromaticity, and other π-systems. Also, condensed results of deeper sequence-based predictions are provided, including secondary structure (total, helical, extended), solvent accessibility (buried), and structural disorder. This may serve as the reference point for a particular case study.

### Charge Decoration Analysis

The Charge Decoration application is designed for interactive comparison of various charge decoration parameters for all LLPS sequences and extracted LLPS disordered fragments. Specifically, the descriptors include sequence charge decoration (SCD)[58], overall charge symmetry (OCS)[59], and a fraction of charged residues (FCR). These recently proposed parameters emerged as a measure of charge distribution along the protein sequence, which, in addition to the overall charge content, turned out as an important factor shaping the protein conformation, especially within low-complexity regions[40]. It is well established that the electrostatic interactions affect the solubility and stabilize the binding interface in some cases of liquid-liquid demixing. In the app, the users can investigate the overall charge decoration patterns and follow the correlations for a complete set of LLPS proteins. Since these measures are a single value per sequence, the background created by the distribution of values for a set of sequences becomes a good reference base in the analysis.

### Statistical analysis of Short Linear Motifs

SLiMs[62] have some unique properties and roles, such as evolutionary conservation, high structural flexibility, involvement in regulatory processes through the intermolecular interactions which makes them relevant for LLPS. The application combines three main tabs delivering the sequence-based detection of (i) eukaryotic linear motifs (ELMs)[34], (ii) other known short stretches of protein sequence such as LARKS[35], Glycine-Arginine Rich regions (GARs)[36], steric zippers[36], or phosphosite motifs[37], (iii) novel motifs filtered out as common from the systematic analysis of various n-mers. The implemented algorithms use the list of motif instances grouped and encoded by regular expressions as keys to search the known phase-separating sequences. The interactive charts allow the user to choose between classes of motifs and show both the total counts and the occurrence in the individual LLPS sequence. The heat map, accessible in the common n-mers section, allows users to track the frequency of specific instances of the 3-mers according to their amino acid composition. In-depth analysis of the user-selected 3-peptide and their most common elongation up to the 6-mers may bring fresh insights about the existence of novel short linear stretches.

### Local Fragments Analysis

While a rough statistical analysis of the full-length sequences, described by a set of averages or fractions, gives general insight, it does conceal some local specificities, which collectively can affect the interaction interface. Sometimes, such flattened characteristics, normalized to the sequence length, can make certain attributes utterly invisible. For example, when reporting the overall density of charge, there is nothing about its distribution along the sequence. Similarly, the frequency of individual amino acids (AA) has no information about their arrangement in particular, whether blocks rich in given AA are present. In light of the recent reports, both properties appear to play crucial roles in the process of LLPS. The Local Fragments category combines the results from a comprehensive analysis conducted on 20-residue fragments extracted along the LLPS sequences with the 5 residues shift. For this purpose, the full-length LLPS sequences (253), structurally disordered LLPS regions (568), and sequences from the SwissProt set (561176, used as a reference benchmark) were fragmented. For these fragments, we examined (i) the distribution of charges (total, positive, negative), (ii) and amino acid compositional diversity, (iii) indicated AA-rich local repeats. The central aim of this analysis is to both identify key residues relevant for phase separation as well as to learn the sequence patterns linked with the LLPS.

### Repository of high-performance pre-computed data

The Download tab, https://biapss.chem.iastate.edu/download.html, provides easy access to our pre-computing bioinformatics resources. Its main aim is to store compressed and ordered data files in a straightforward CSV format giving users access to high-quality sequence-based results from analysis coupled with additional features, such as rich annotations to external databases and the agreed consensus of predictive methods. The available data includes raw predictions which are pre-calculated using a set of well-established tools as well as findings of our deep statistical analysis of overall characteristics of LLPS sequences. For the user’s convenience, we have unified and integrated the results from these sources into several categories following the list of applications in the SingleSEQ and MultiSEQ tabs. We have attached descriptions for each file to standardize the data and make the content easier to understand.

## Availability and future directions

### Software availability and requirements

BIAPSS is a free online resource accessible at https://biapss.chem.iastate.edu/. The web interface of BIAPSS is developed within the Flask/Python web framework using HTML, CSS, and Plotly-Dash graphing libraries, with all the major browsers supported including the mobile device accessibility. Suggested browsers include:

- for Linux/Ubuntu users: Google Chrome, Firefox
- for Windows users: Google Chrome, Firefox, Opera
- for Mac users: Safari
- for mobile devices: the website is adapted to mobile devices larger than 600×300px and some of its functionality may be limited.

### Documentation

The comprehensive documentation for the BIAPSS services is available online (https://biapss.chem.iastate.edu/documentation.html). The provided documentation gives detailed instructions on how to browse the single LLPS sequence analysis results and how to interpret the statistics of the complete LLPS dataset. The user can explore selected sections and learn more about methods used from the literature sources listed in the Tools & References. For the new users, BIAPSS provides many simple tutorials demonstrating the usage of available web applications highlighting their options and features. Specifically, the orange *Quick Guide* button is located in the top-left corner of the particular web application. Finally, the detailed user manual is provided as a PDF file in the Supporting Information of this manuscript.

### Improvements and community outreach

Future efforts will be made to keep the repository current by regularly updating the repository. We are currently working on several new applications for other potentially LLPS-relevant factors, including aggregation and prion-like propensity, detailed π-π contact frequency, and more. Future plans include adding a section dedicated to the structure-based statistics and visualization for cases with known fragments of three-dimensional structure, both determined experimentally or projected in comparative modeling. Furthermore, since potential users may have their own sequences of interest, either natural or designed, we plan to create an upload section to parse user-defined cases and compare them with the benchmark of known LLPS-driver proteins. Overall, when the BIAPSS web platform grows in size and accuracy, we will take up the challenge of developing a robust predictor detecting regions relevant for phase separation or suggest the sequence modifications that could modulate the phase behavior.

## Supporting Information

Supporting_Information.pdf - the file contains an extensive user manual for the BIAPSS web platform.

## Author Contribution

Conceptualization, A.E.B-D.; Software development, A.E.B-D.; Writing an original draft, A.E.B-D. and D.A.P.

## Acknowledgments

A.E.B-D. acknowledges generous financial support by Roy J. Carver Charitable Trust through Iowa State University Bioscience Innovation Postdoctoral Fellowship. This work was supported by the National Institute Of General Medical Sciences of the National Institutes of Health [R35GM138243 to D.A.P.]. The content is solely the responsibility of the authors and does not necessarily represent the official views of the NationalInstitutes of Health.

