## Supporting Information for "BIAPSS - BioInformatic Analysis of liquid-liquid Phase-Separating protein Sequences"

SUPPLEMENTARY INFORMATION

### BIAPSS: BIoinformatic Analysis of LLPS Sequences

availability: *https://biapss.chem.iastate.edu/*

---

#### The layout of BIAPSS web platform

BIAPSS is designed as a user-friendly web platform for interactive sequence-based analysis of protein which are experimentally confirmed to undergo liquid-liquid phase separation. The layout and main functionalities of BIAPSS services are summarized in the Figure S1. The web platform provides (i) a database of the superset of experimentally evidenced LLPS-driver protein sequences, (ii) a repository of pre-computed data of comprehensive bioinformatics statistics (available via *DOWNLOAD* tab), (iii) two sets of web server applications supporting the interactive analysis and visualization of physicochemical characteristics (available via *SingleSEQ* and *MultiSEQ* tabs). All graphs generated by the BIAPSS web applications are based on the Plotly-Dash libraries for interactive graphing. Broad documentation, containing details of the employed methods and the external tools, are provided in the *DOCS* tab.

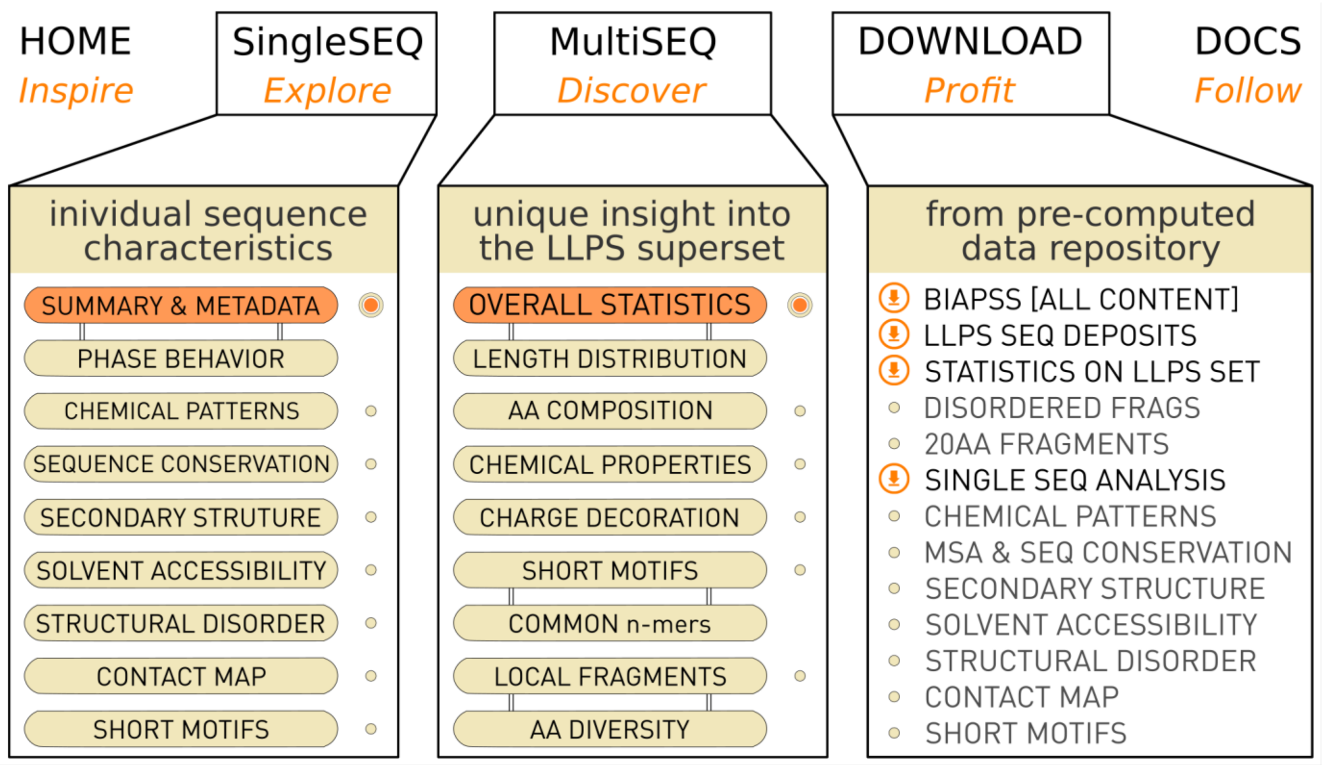


Figure S1. The layout of BIAPSS web platform (https://biapss.chem.iastate.edu) for comprehensive sequence-based analysis of LLPS proteins. The core of the implemented web applications and data repository is contained in the SingleSEQ, MultiSEQ, and Download tabs.

#### SingleSEQ TAB - Applications for an individual sequence characteristics

availibility: *https://biapss.chem.iastate.edu/single_seq.html*

---

##### *Summary & Metadata App*

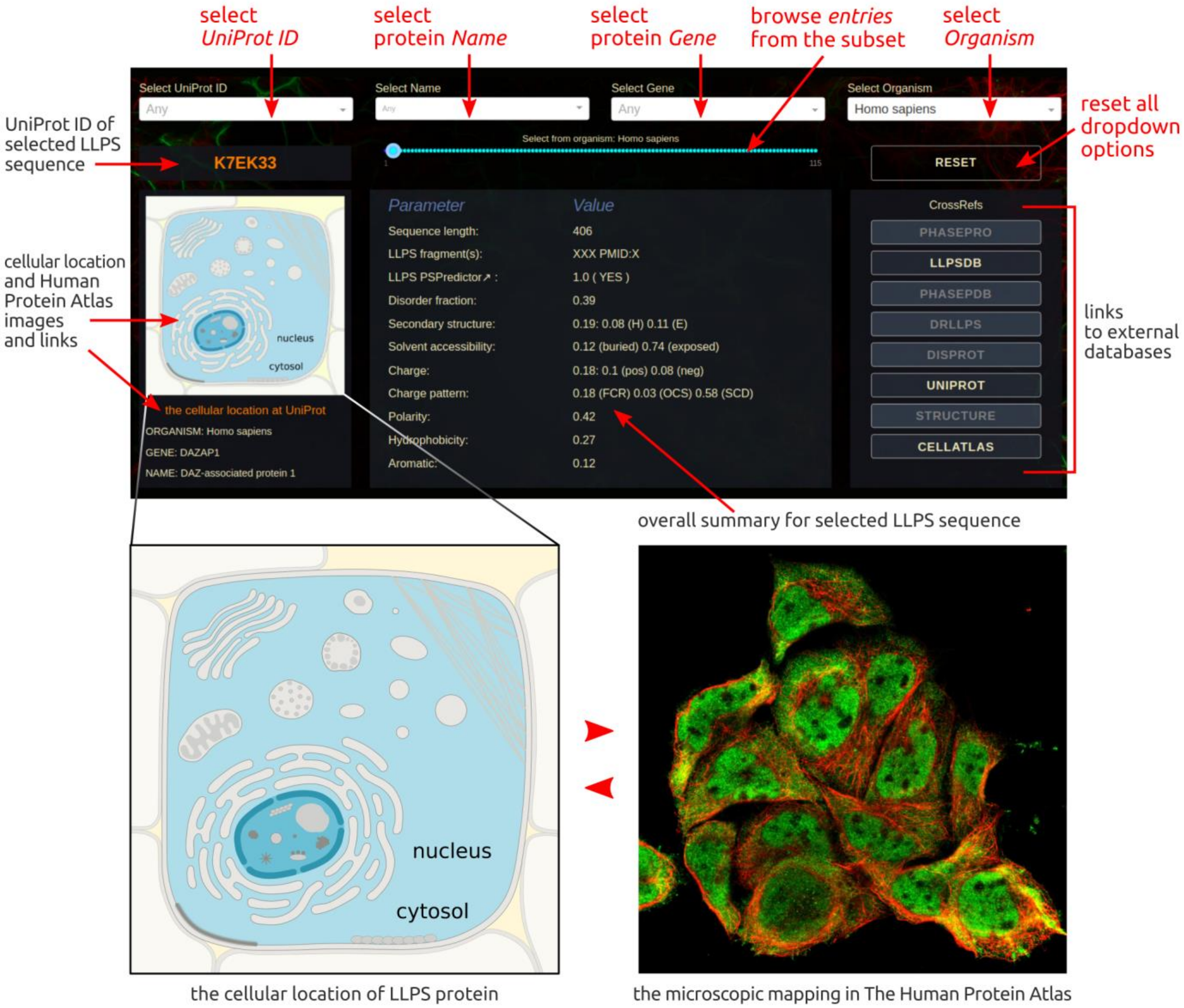


Figure S2. Snapshot of the Summary & Metadata web application with the main functionalities highlighted with the red arrows.

The Analysis of Single LLPS Sequence page provides a compact one-page summary for a single LLPS protein sequence. The sequence can be selected individually from the list of BIAPSS deposits using the 2-row querying panel on the top. The first row contains four dropdown menus, which work as a filter narrowing the searched subset of entries. For user convenience, the **dropdowns are labeled** on the top and are searchable, i.e., they limit a subset of results to the keyword entered by the user. Looking from the left, the following **option boxes** allow querying the database by (i) UniProt identifier, (ii) common protein name, (iii) corresponding gene, (iv) organism. Accordingly, in the second row, the **horizontal slider** in the middle (cyan) shows the size of a reduced subset, e.g., to a chosen *Homo Sapiens* organism and subsequent interactive dots load the content for each entry. The **reset button** on the right lets the user return to the default settings and start a new search, while the orange text field on the left gives the UniProt identifier for the currently displayed content.

The data section in this application consists of 3 vertical panels. On the left, the user can find basic information about the sequence, such as a common **name**, **organism** of origin, **gene**, and a graphic illustration of the **subcellular location**, as the consensus of data derived from UniProt, COMPARTMENTS, and available primary LLPS databases. For human proteins, swapping with cellular localization, a **microscopic image** of the cell is embedded via The Human Protein Atlas. The tooltip below the image provides not only the list of sources for the data but also links directly to them. In the central panel, the user can find a compact summary of the key **biophysical properties and sequence-based characteristics** for a given LLPS protein sequence. Among the parameters, the user can find the **sequence length**, **LLPS-related region(s)** together with corresponding **literature PMID**, PSPredictor scores estimating the **propensity to phase behavior**, calculated **fractions of charged, polar, hydrophobic, and aromatic residues**, and values of various **charge decoration** parameters (OCS, SCD, FCR). Also, the predicted fractions of the following: **structurally disordered residues**, **secondary structure** (helical/extended residues), **solvent accessibility** (buried/exposed residues) are provided. All values are consensus averages that were derived from the sequence-based predictions of several state-of-the-art methods. The right-side Cross-References section combines multiple external sources that provide **relevant annotations** complementary to the BIAPSS resources. These annotations are the **primary LLPS-related databases**, such as PhaSepDB, drLLPS, LLPSDB, and PhaSePro; **experimental structure** (if known, then linked to the BDBE-KB repository, otherwise the computationally modeled structure is accessible via the MODBASE database and/or SWISS-MODEL server); manually curated **intrinsically disordered regions** in DisProt database and **protein sequence** entry in the UniProt database. The one-click buttons links to the corresponding entry in the related databases.

#### *Chemical Patterns App*

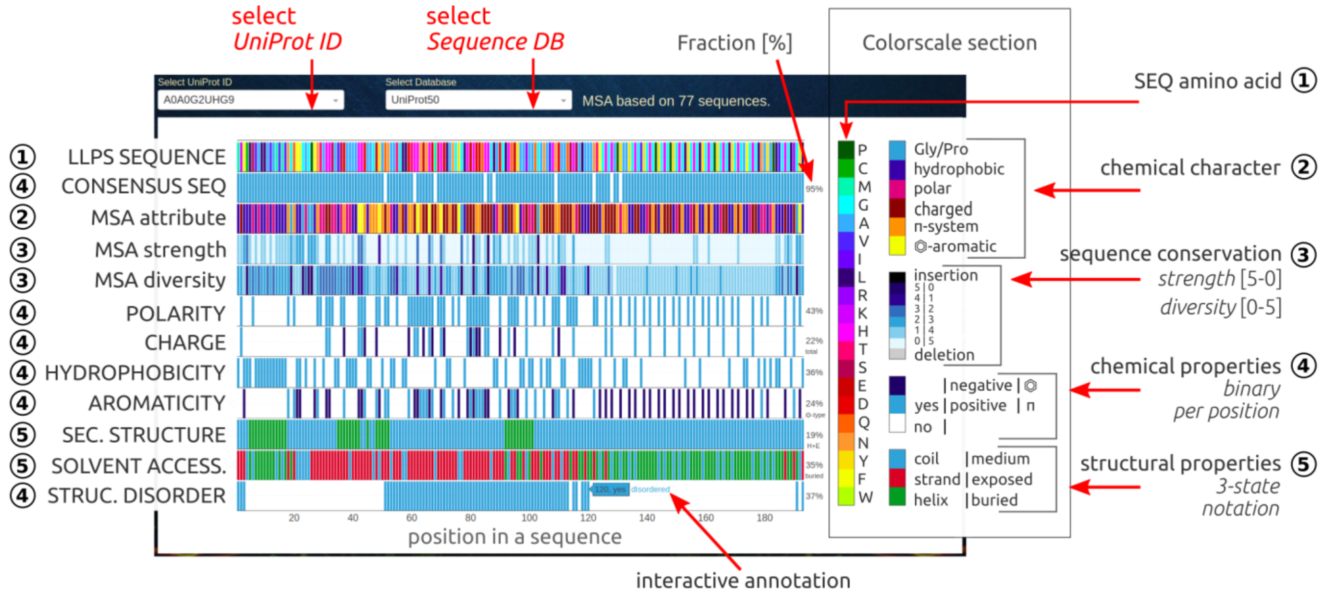


Figure S3. Snapshot of the Chemical Patterns web application with its main functionalities highlighted by red arrows.

The Chemical Patterns application allows comparing all key properties for a given single amino acids sequence all within a single chart. From the **consensus sequence of multiple sequence alignment** (MSA) the **evolutionary conservation** (including **insertions** and **deletions**) can be identified as the regions which indicate functional or structural similarity. Three additional MSA-based parameters describe per position **attribute** - the character of most common amino acids, **diversity** - the number of various amino acids, **strength** - the conservation level. The chemical properties such as **polarity**, **hydrophobicity**, **aromaticity** as well as **charge** pattern are provided. The consensus of several state-of-the-art methods for prediction of **secondary structure** (SS3), **solvent accessibility** (SA) and **structural disorder** are pre-calculated. For each parameter, the fraction of residues in the sequence that meets the criterion is also given. Overview of all parameters in one place helps to identify key residues or short regions that can play a crucial role in LLPS behavior or other functional mechanisms. Each position in the sequence is annotated interactively.

##### *Dropdown options*

The user can select the LLPS sequence by its UniProtID. The second dropdown allows selecting the subset of UniProt sequences (among the smallest SwissProt, moderate UniRef50, or the largest UniRef90 databases) against which the multiple sequence alignment (MSA) was calculated.

##### *Graph section*

*1)* The first row is the **input sequence** colored by the amino acid composition. The interactive annotation per position in the sequence includes residue index and the amino acid type.

*2)* The second row is the **consensus sequence** obtained from the profile of multiple sequence alignment. The positions being identical with the query sequence are marked in blue, the positions differing in white. The interactive annotation per sequence position includes the residue index and amino acid type. The number on the right indicates the percentage identity of the consensus and the query sequence. The consensus sequence can vary depending on the selected reference database (SwissProt, UniRef50, UniRef90), and number of related sequences used in the MSA calculation (the result is reliable if at least several dozen sequences have been used).

*3)* The **MSA attribute** describes the chemical nature of the most common amino acids at a given position in the multiple sequence alignment. The defined attributes include: polar (magenta), charge (dark-red), π-system (orange), aromatic (yellow), hydrophobic (navy), other (G or P, blue).

*4)* The **MSA diversity** define the number of various amino acids detected at a given position in the alignment - the value of this parameter is a discrete number in the range 0-5 (0 - single AA, 1 - two AA, 2 - three AA, 3 - four AA, 4 - five or six AA, 5 – 7 and more AA). The interactive annotation includes an index at the sequence, diversity value, and list of corresponding amino acid types at a given position in the alignment.

*5)* The **MSA strength** is a parameter describing the degree of conservation (in the range 0-5: 0 - not conserved, 5 - highly conserved) derived from the results of *hmmlogo* (included in HMMER3.3 package) calculations.

Attribute, strength, and diversity are the newly designed (for the purpose of this work) three parameters related to MSA calculation that characterize the evolutionary conservation of the query sequence. Diversity and strength parameters use the same discrete range while reversed color scales, so the visual interpretations of these two rows is as simple as the meaning of the darkest bars (navy) that correspond to the highest conservation and lowest diversity (conserved position in alignment and type of amino acid). The quick analysis of such position, especially when the query sequence at a given point is different (i.e., consensus bar is white) or deleted (sequence bar is gray), helps to detect the sensitive regions in the sequence that play a key role in the interactions important for structural stability or biological activity.

*6)* The sixth row contains a binary pattern (blue-white) of **polar residues** (S, D, N, T, E, Q, H, K, R, Y) in the query sequence. The number on the right indicates the percentage content of polar residues.

*7)* The seventh row contains a ternary pattern (blue-navy-white) of **charged residues** in the query sequence: positive charge (H, K, R) is marked in blue, while the negative charge (D, E) is marked in navy. The number on the right indicates the percentage content of all charged residues.

*8)* The eighth row contains a binary pattern (blue-white) of **hydrophobic residues** (A, C, V, M, I, L, F, W) in the query sequence. The number on the right indicates the percentage content of hydrophobic residues.

*9)* The ninth row contains the ternary pattern (blue-navy-white) of **aromatic residues** (W, F, Y, H) marked in navy and other π-system containing residues (R, N, Q, D, E) marked in blue. The number on the right indicates the percentage content of aromatic residues.

*10)* The tenth row contains the consensus of **secondary structure** assignment provided in 3-state notation (H - helical, marked in green; E - extended, marked in red; C - coil, marked in blue) and predicted by several state-of-the-art methods (PSIPRED, RAPTOR-X, PORTER-5, SPIDER-3, FESS). The number on the right indicates the percentage of content residues that are assigned to the secondary structure element (helix or strand).

*11)* The eleventh row contains the consensus of **solvent accessibility** assignment provided in 3-state notation (B - buried, marked in green; E - exposed, marked in red; M - medium, marked in blue) and predicted by several state-of-the-art methods (RAPTOR-X, PaleAle5, SPOT-1D). The number on the right indicates the percentage of content of buried residues.

*12)* The twelfth row contains the consensus of **structural disorder** (DISO) predicted by several state-of-the-art methods (RAPTOR-X, IUPred2A, SPOT-Disorder, DISOPRED2, DISOPRED3, VSL2, PONDR-FIT, PONDR-VLXT). The number on the right indicates the percentage content of disordered residues.

#### ***Sequence Conservation App***

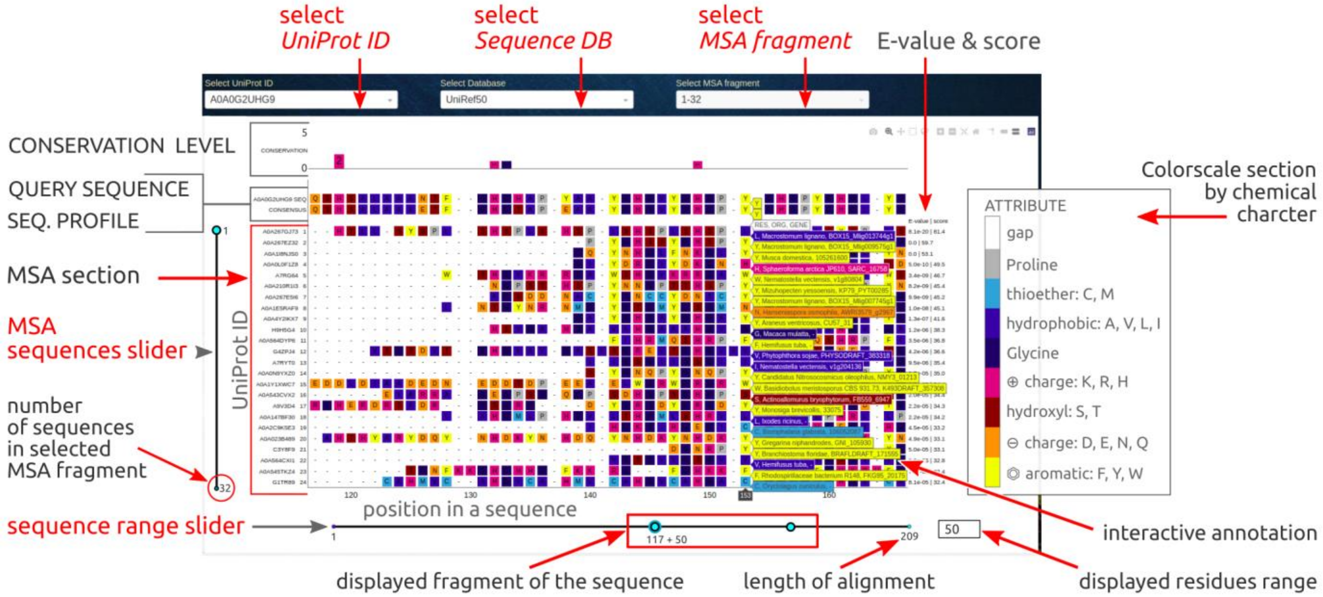

Figure S4. Snapshot of the Sequence Conservation web application with its main functionalities highlighted by red arrows.

The Sequence Conservation application contains the **Multiple Sequence Alignment** (MSA) of LLPS sequences against sequences selected from the UniProt subsets (varied in size and sequence identity). The MSA, **consensus profile**, and simplified **sequence conservation** are assigned per position in the query LLPS-sequence. The conservation strength may be a consequence of the functional, structural, or evolutionary importance of the region in the sequence. Therefore, the unique **insertions**, **deletions**, or **substitutions**, identified within the LLPS protein sequence can help to identify the regions relevant to the formation of biological condensates.

A multiple sequence alignment (MSA) is a sequence alignment of three or more biological sequences - here we analyzed the protein sequences composed of amino acids. Alignment is matching different sequences from regions which have more similarity with each other while the multiple sequence alignment tries to minimize the number of gaps (deletions/insertions) and produce a compact alignment that can be used to build the consensus profile (amino acids at each position in the alignment are scored according to the frequency with which they occur). The position-specific scoring systems allow determining the evolutionary conservation of protein sequences at the level of individual amino acids. The MSA available in the web application was prepared by using an efficient HMMER3.3 method that employs a probabilistic hidden Markov model (HMM). HMMER makes a probabilistic profile of the query sequence that assigns a position-specific scoring system for substitutions, insertions, and deletions. Compared to BLAST-based searches, that method is significantly more accurate (detect also the remote homologs) and similarly fast for protein search.

##### *Options (dropdowns and sliders)*

The user can select the LLPS sequence by its UniProtID from the first dropdown. The second dropdown allows selecting the subset of UniProt sequences (among the smallest SwissProt, moderate UniRef50, or the largest UniRef90 databases) against which the multiple sequence alignment (MSA) was calculated. Since the MSA is usually large, the displayed fragment is controlled on-the-fly by the third dropdown (loads packets of 500 sequences sorted by e-value due to memory optimization) and two sliders (vertical and horizontal) that define (i) the list of the displayed sequences (24 consecutive ones) and (ii) a sequence fragment of a selected length (from 10 to 500 residues, step=5) along the profile.

#### *Graph section*

***1) The sequence conservation section***

The conservation level (the upper section) in the discrete range from 0 to 5 was assigned per position based on results returned from HMMER3.3 *hmmlogo* tool. This section is annotated by consensus (profile) sequence and the most common amino acid variants at the position.

***2) The sequence profile section***

The consensus profile was prepared by using the efficient HMMER3.3 method (*hmmbuild*) that employs a probabilistic hidden Markov model (HMM) with setting position-specific gap and insertion scores. The profile captures important information about the degree of conservation at various positions in the multiple alignments. The used colors correspond to the chemical character of amino acid: yellow - aromatic (F, Y, W), orange - acidic or negatively charged (D, E, N, Q), dark-red - hydroxyl-containing (S, T), magenta - positively charged (K, R, H), navy - glycine, purple - hydrophobic (A, V, L, I), sulfur-containing (C, M), gray - proline, white - gap.

***3) Multiple Sequence Alignment section***

MSA section contains aligned protein sequences colored by the chemical character of amino acid and sorted by E-value (score is also provided). Each position in the sequence is annotated interactively by index at alignment profile, amino acid, organism, and gene. The displayed fragment of the MSA is controlled on-the-fly by two sliders. The multiple sequence alignment was produced by using the HMMER3.3 method (*phmmer* + *hmmalign*) to search the selected sequence database (SwissProt, UniRef50, UniRef90).

The sequence **E-value** is the measure of statistical significance, i.e., the expectation of finding that sequence by random chance or the expected number of false positives (nonhomologous sequences). The lower the E-value, the more significant the hit (here, the sequences with ≤ 10−3 or so were considered to be significant hits). The HMMER E-value is based on the sequence bit score, which is the log-odds score for the complete sequence and doesn't depend on the size of the sequence database (only on the profile and the target sequence).

#### Secondary Structure App

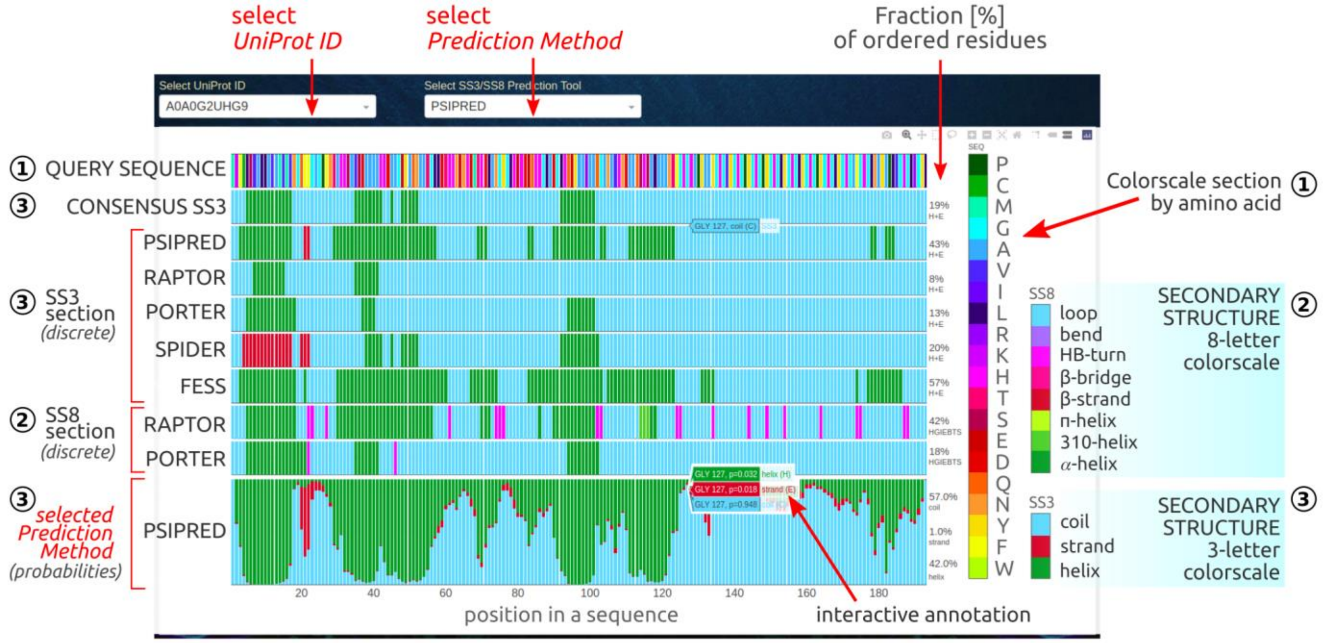


Figure S5. Snapshot of the Secondary Structure web application with its main functionalities highlighted by red arrows.

The Secondary Structure application presents the sequence-based predictions of **secondary structure** elements assigned along the selected LLPS sequence. The user can explore with residue-resolution (i) the **consensus assignment** in SS3 notation, (ii) the **original SS3/SS8 assignments** obtained from state-of-the-art methods, and (iii) **detailed probabilities** of helical, extended or coiled structure for individual amino acids along the LLPS sequence. Also, the column on the right provides the **total fraction of ordered residues**.

The secondary structure is a spatial arrangement of short, containing up to 30 consecutive amino acid fragments of the polypeptide chain into one of the known ordered structure elements: **helical** (α, $3_{10}$, π), **extended** (β-strand, β-bridge, HB-turn, bend), or disordered (**coil,** loop). The appearance of regions taking a strictly defined shape in space is conditioned by the presence of a series of periodically repeatable values of *Phi* and *Psi* angles in the protein main chain. It is a sequence of amino acids forcing the pattern of hydrogen bonds between backbone atoms that determine the formation of spatially and energetically stable conformation of a particular secondary structure. Due to the local character of the geometric order of the protein backbone, various elements of the secondary structure may be present in a single polypeptide chain. The recent findings suggest that some intrinsically disordered proteins can temporarily form an orderly structure when the conditions are favorable, e.g., when a binding partner provides energetic stability. Therefore, the predicted probabilities to form secondary structure elements opens up the possibility to detect sequence fragments slightly biased to form an ordered structure, such as steric zipper or LARKS, and provides a better understanding of the role that the local amino acid composition plays in supporting multivalent interactions.

##### Dropdown options

The user can select the LLPS sequence by its UniProtID. The second dropdown allows selecting the prediction method used to calculate the detailed probabilities of various secondary structure elements.

#### *Graph section*

*1)* The first row is the **input sequence** colored by amino acid composition. The interactive annotation per position at sequence includes residue index and amino acid type.

*2)* The second row is the **consensus secondary structure** assignment provided in 3-letter notation (SS3: **H** - helix, **E** - strand, **C** - coil), derived as the agreed outcome of 5 well-established methods (PSIPRED, RAPTOR-X, PORTER-5, SPIDER-3, FESS).

*3) SS3 section – secondary structure in 3-letter notation (HEC)*

The rows 3-7 are the original predictions of secondary structure obtained from the benchmarked methods, and provided as binary residue-resolution assignment in **SS3 notation** along the protein sequence. The used colors encode: cyan – helix (**H**), red – extended (**E**), green – helix (**C**).

*4) SS8 section – secondary structure in 8-letter notation (HGIEBTSC)*

The rows 8 and 9 are the original predictions of secondary structure obtained from two the most advanced methods (RAPTOR-X and PORTER-5), and provided as binary residue-resolution assignment in **SS8 notation** along the protein sequence. The used colors encode: forest – α-helix (**H**), green – $3_{10}$-helix (**G**), lime – π-helix (**I**), red – β-strand (**E**), magenta – β-bridge (**B**), pink – HB-turn (**T**), purple – bend (**S**), cyan – loop (**C**).

*5) Probabilities section*

For a selected method, in the probabilities section, the **detailed fractions of each type of secondary structure** elements are provided in SS3 notation per position in the protein sequence. The interactive annotations include residue index, amino acid type and probabilities for HEC elements.

#### *Solvent Accessibility App*

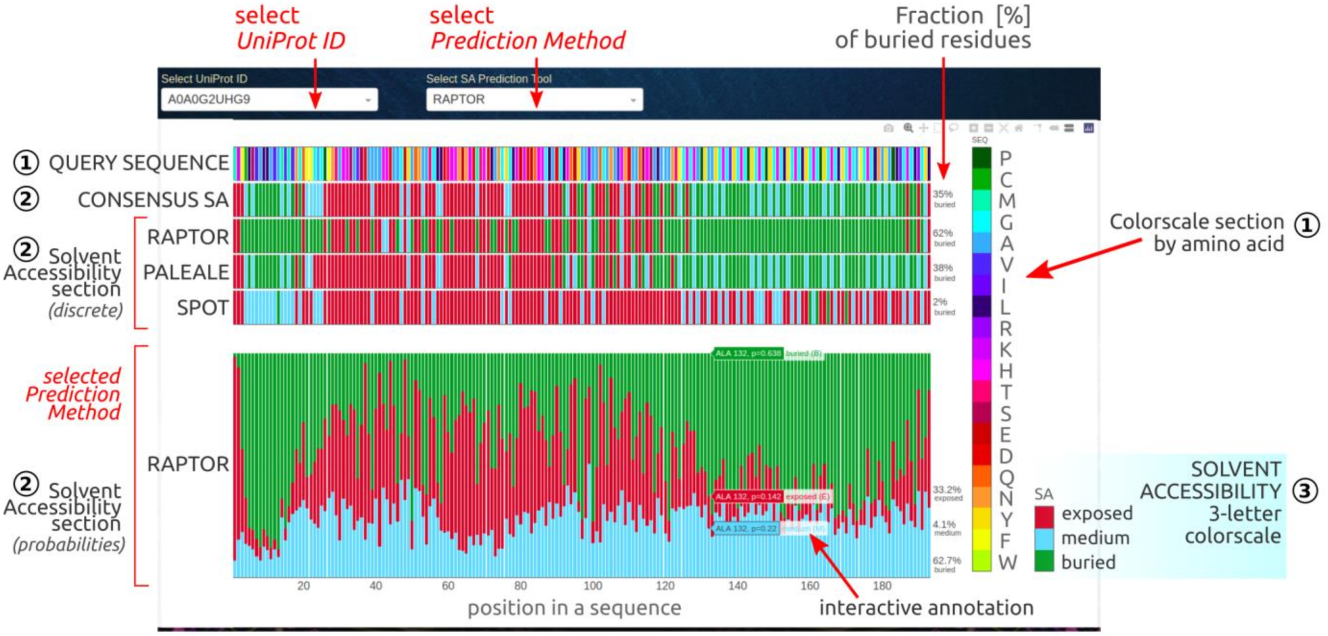


Figure S6. Snapshot of the Solvent Accessibility web application with its main functionalities highlighted by red arrows.

The Solvent Accessibility application presents the sequence-based predictions of the tendency of residues to be **buried** in the interior of the protein or to be **exposed** to the solvent. The user can explore with residue-resolution (i) the **consensus assignment** of solvent accessibility in 3-state notation (exposed, medium, buried), (ii) the **original assignments** obtained from state-of-the-art methods, and (iii) **detailed probabilities** of exposed or buried surface for individual amino acids along the LLPS sequence. Also, the column on the right provides the **total fraction of buried residues**.

Solvent accessibility is a measure of hydrophobic stabilization, usually defined as the accessible surface area (ASA) or solvent-accessible surface area (SASA) and described in units of square Ångstroms. The solvent accessibility of protein residues is one of the driving forces (hydrophobic effect) of protein folding as it is related to the spatial arrangement and packing of the hydrophobic core of protein chain. Even a rough sequence-based prediction of the solvent accessibility can provide structurally relevant information to identify the binding interface or multivalent regions important for the liquid-liquid phase separation.

##### *Dropdown options*

The user can select the LLPS sequence by its UniProtID. The second dropdown allows selecting the prediction method used to calculate the detailed probabilities of solvent accessibility.

##### *Graph section*

*1)* The first row is the **input sequence** colored by amino acid composition. The interactive annotation per position at sequence includes residue index and amino acid type.

*2)* The second row is the **consensus solvent accessibility** assignment provided in 3-state notation (**E** - epxosed, **B** - buried, **M** - medium), derived as the agreed outcome of 3 well-established methods (RAPTOR-X, PaleAle5, SPOT-1D).

*3) Solvent accessibility section (EBM)*

The rows 3-5 are the original predictions of solvent accessibility obtained from the benchmarked methods, and provided as binary residue-resolution assignment in 3-state notation along the protein sequence. The used colors encode: cyan – medium (**M**), red – exposed (**E**), green – buried (**B**).

*4) Probabilities section*

For a selected method, in the probabilities section, the **detailed fractions of buried and exposed surface** are provided per position in the protein sequence. The interactive annotations include residue index, amino acid type and probabilities for EBM elements.

#### Structural Disorder App

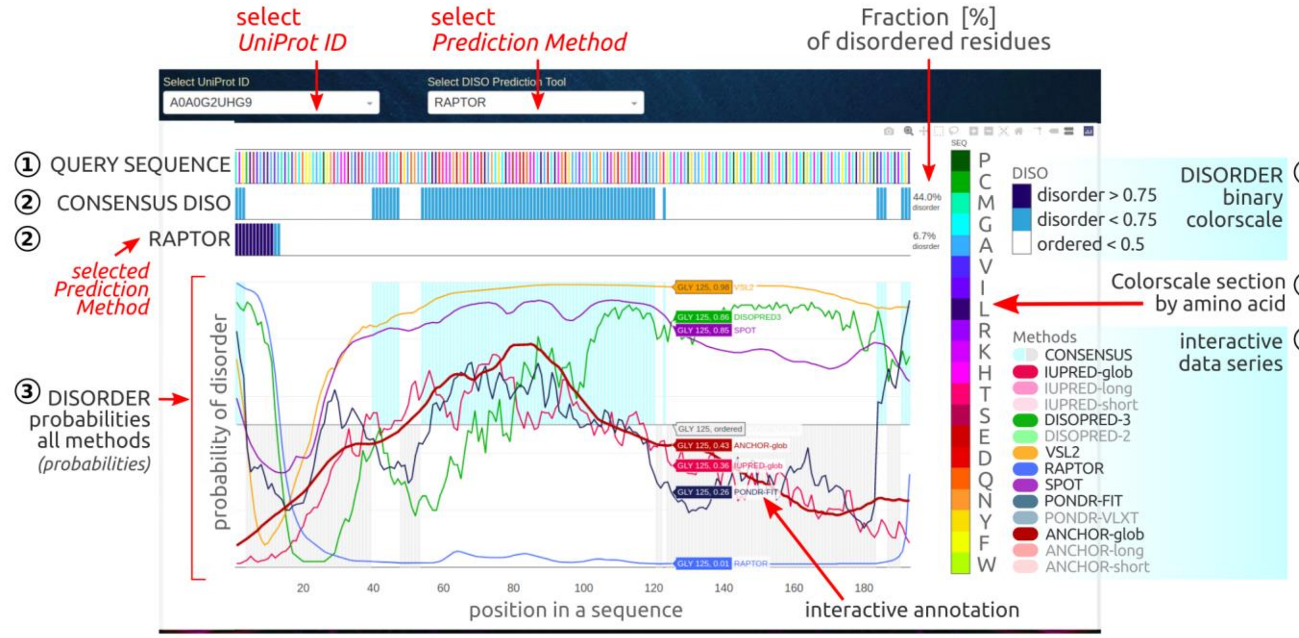


Figure S7. Snapshot of the Structural Disorder web application with its main functionalities highlighted by red arrows.

The Structural Disorder application contains the sequence-based predictions of **disordered regions** provided as binary assignment at a given position in the sequence. The user can explore with residue-resolution (i) the **consensus assignment** of disordered regions, (ii) the **original assignment** obtained from user-selected state-of-the-art method, and (iii) **detailed probabilities** of structural disorder obtained from all the benchmark methods. Also, the column on the right provides the **total fraction of disordered residues**.

Structural disorder of protein is defined by the lack of fixed or ordered three-dimensional structure. The definition covers wide range of cases from fully unstructured to partially structured.

##### *Dropdown options*

The user can select the LLPS sequence by its UniProtID. The second dropdown allows selecting the prediction method and comparing the original results to the agreed outcome from all the benchmark tools.

##### *Graph section*

*1)* The first row is the **input sequence** colored by amino acid composition. The interactive annotation per position at sequence includes residue index and amino acid type.

*2)* The second row is the **consensus structural disorder** provided in binary assignment, derived as the agreed outcome of 7 well-established methods (RAPTOR-X, IUPred2A, SPOT-Disorder, DISOPRED2, DISOPRED3, VSL2, PONDR-FIT, PONDR-VLXT).

*3)* The third row is the original prediction of **disordered regions obtained from the user-selected method**, and provided as binary residue-resolution assignment along the protein sequence. The used colors encode: navy – definitely disordered (p > 0.75), blue – fairly disordered (0.5 < p < 0.75), white – unlikely disordered (p < 0.5).

*4) Probabilities section*

For all the methods, in the probabilities section, the **detailed probability of disorder** is provided per position in the protein sequence. The responsive data series on the right allows displaying customized results. The interactive annotations include residue index, amino acid type and and disorder probability value. The ANCHOR method was used for predicting **protein binding regions** in disordered fragments.

#### *Contact Map App*

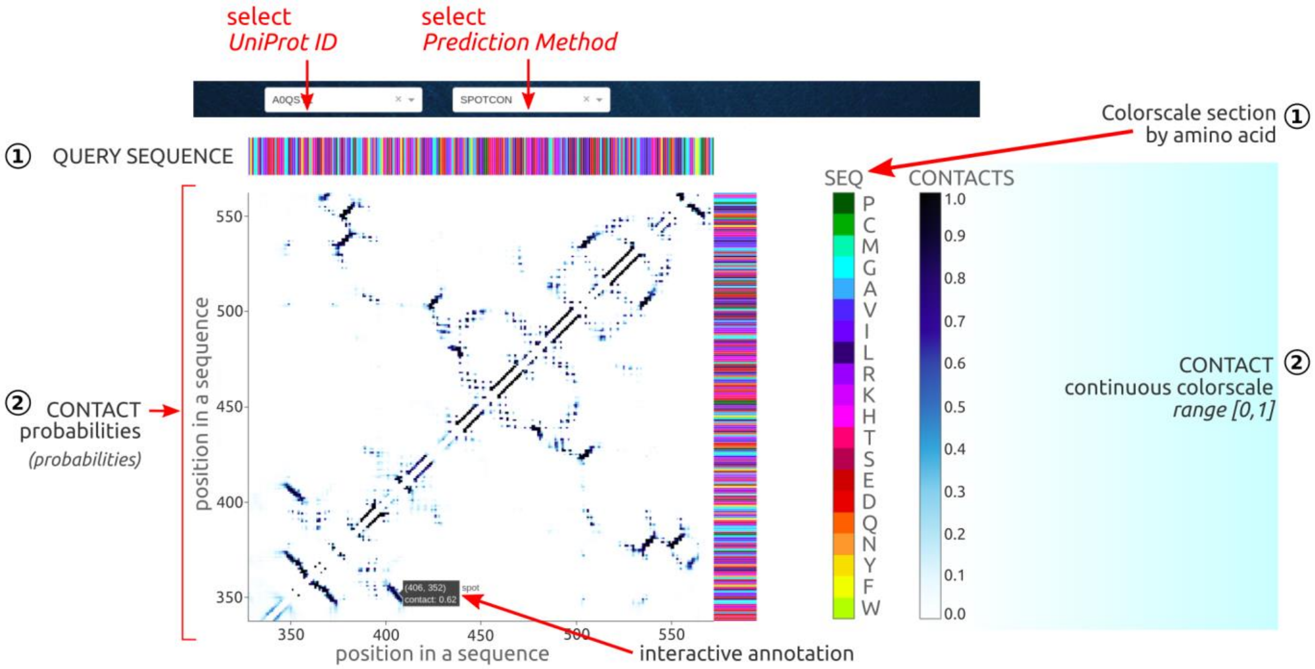


Figure S8. Snapshot of the Contact Map web application with its main functionalities highlighted by red arrows.

The Contact Map application contains the sequence-based predictions of **spatial contacts** between residues in protein sequence provided as square heatmap of **contact probabilities**. The pair of residues is considered as being in spatial contact for probability value above 0.5. An in-depth analysis of the contacts predicted for LLPS proteins, even though they are largely disordered, can help to identify regions essential for the maintaining structure, dynamics, function, and may indicate a lead relevant to phase behavior.

Contact maps provide a more reduced representation of a protein structure using a binary two-dimensional matrix of distances between all possible amino acid residue pairs. The contact number of protein residues limits the possibilities of protein conformations and helps to encode a three-dimensional structure. The sequence-based prediction of contacts is feasible with the availability of high numbers of sequences that contain coupled (coevolving) residue pairs. The most powerful contact predictors, such as selected for this work Raptor-X Contact, SPOT-Contact, and ResPRE methods employ the convolutional neural networks, machine learning and evolutionary coupling techniques.

##### *Dropdown options*

The user can select the LLPS sequence by its UniProtID. The second dropdown allows selecting the contact map predicted by one of 3 well-established methods: RATOR-X Contact, SPOT-Contact, ResPRE.

##### *Graph section*

*1)* The top row and right column are the **input sequence** colored by amino acid composition. The interactive annotation per position at sequence includes residue index and amino acid type.

*2)* The square heatmap presents the **spatial contacts** predicted by user-selected method. The interactive annotation shows the detailed probability of contact between the pair of interacting residues.

#### *Short Linear Motifs App*

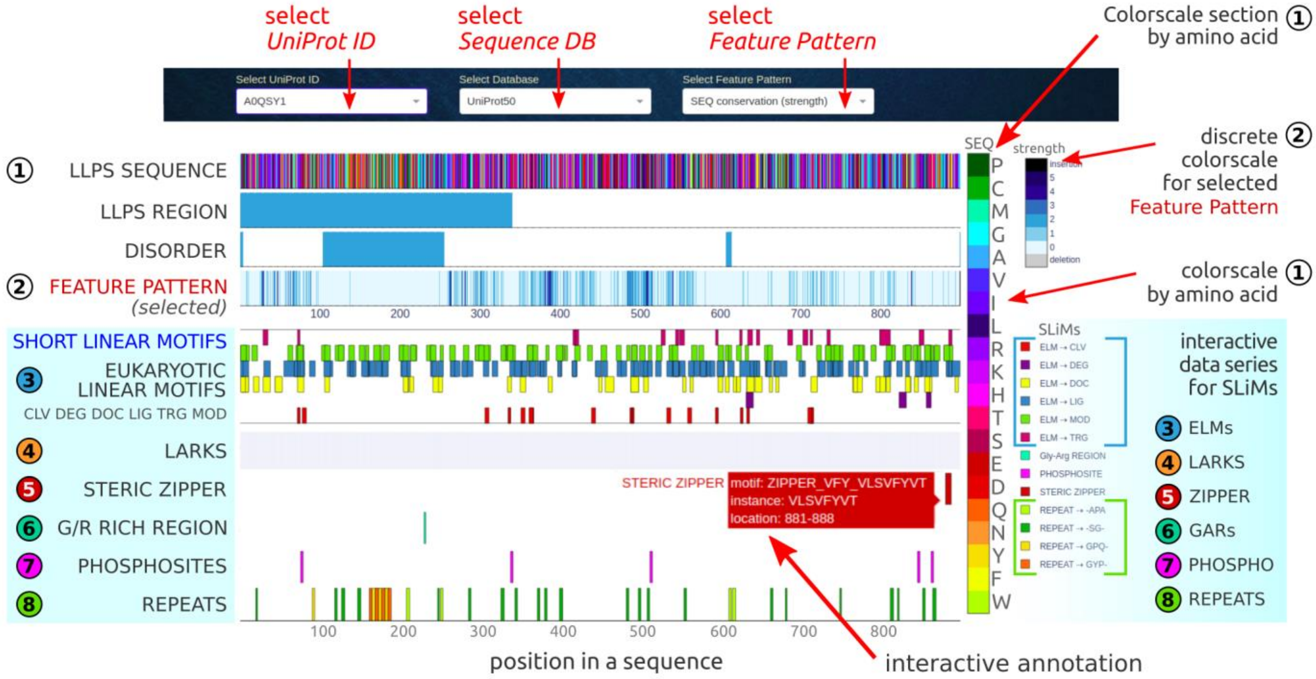


Figure S9. Snapshot of the Short Linear Motifs web application with its main functionalities highlighted by red arrows.

The Short Motifs SEQ application is designed to detect various **short linear motifs** (SLiMs) in the individual LLPS sequence in reference to regions experimentally evidenced as LLPS-related and/or were predicted to be structurally disordered. The evolutionary conservation of SliMs within intrinsically disordered fragments, clearly points to their significance for the regulatory function, hence may also be essential for the collective formation of membraneless organelles. The multi-row application offers a broad screening of short motifs known from literature as relevant for phase behavior, such as the low-complexity aromatic-rich kinked segments (**LARKS**), glycine-arginine rich regions (**GARs**), **steric zippers** and S--D/S--E motifs specific for **phosphosites**. Accordingly, the eukaryotic linear motifs (**ELMs**), experimentally validated in eukaryotic cells, are indicated along the sequence with a discrimination of the main ELM classes. Additionally, short fragments frequently repeated along an individual sequence are detectable, including random and tandem repeats.

##### *Dropdown options*

The user can select the LLPS sequence by its UniProtID. The second dropdown allows selecting the reference sequence database used in multiple sequence alignments and sequence-based predictions of various structurally-relevant properties. The third dropdown allows for incorporating various datasets that serve as a reference background and extend the available analysis to user-specific needs. Customization includes loading any sequence-based characteristics available in other applications as a single-row result.

##### *Graph section*

*1)* The first row is the **input sequence** colored by amino acid composition. The interactive annotation per position at sequence includes residue index and amino acid type.

*2)* The second row is the **sequence region** experimentally confirmed to be **involved in protein phase separation**. The interactive annotation includes residue index, amino acid type and PubMed identifier to corresponding literature.

*3)* The third row is the **consensus structural disorder**, derived as the agreed outcome of 7 well-established methods (RAPTOR-X, IUPred2A, SPOT-Disorder, DISOPRED2, DISOPRED3, VSL2, PONDR-FIT, PONDR-VLXT).

*4)* The fourth row is a single-row **characteristic selected by the users** to support their analyses. The specific feature pattern can be customized via third dropdown (see available options in the Figure S9).

*5) Short linear motifs section*

The short linear motifs section detects short stretches along the protein sequence, divided on six main groups:
• **ELMs**, the Eukaryotic Linear Motifs experimentally validated in eukaryotic cells;
• **LARKS**, the low-complexity aromatic-rich kinked segments;
• **steric zipper**, a structural motif of dry interface formed via van der Waals interactions and hydrogen bonds;
• **GARs**, Glycine-Arginine Rich Regions;
• **PTMs**, the post-translational modification sites (S--D/S--E motifs for phosphosites);
• **repeats**, short fragments often repeated along the sequence.

The detailed information (location, motif instance, value) is displayed on interactive labels when hovering over a point in the chart. The motif’s location can be correlated to main sequence's characteristics provided in the top four app's rows.

#### MultiSEQ TAB - Statistics on the superset of LLPS Sequences

availability: *https://biapss.chem.iastate.edu/statistics.html*

---

##### *Overall Statistics App*

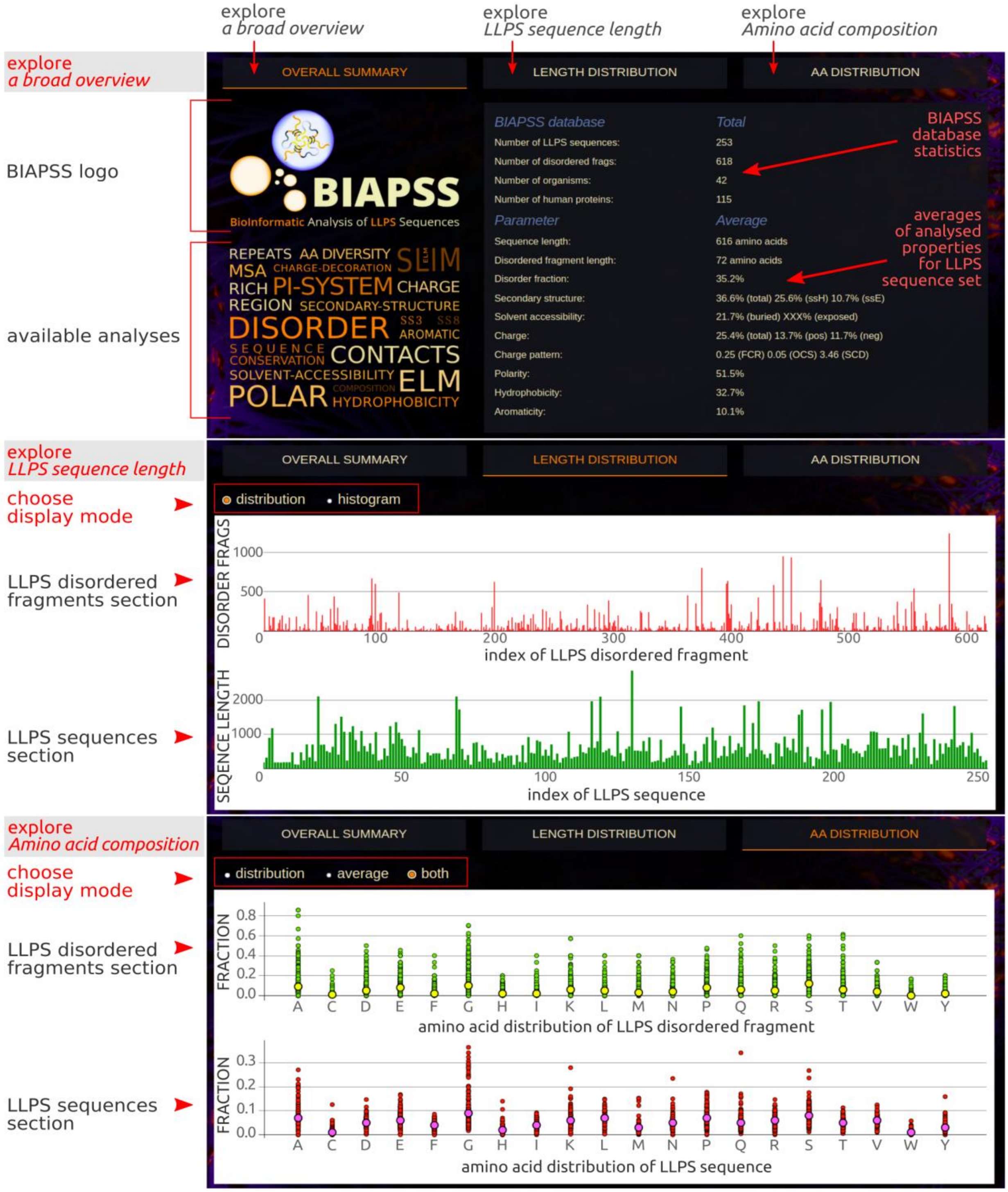


Figure S10. Snapshot of the Overall Statistics web application with its main functionalities highlighted by red arrows.

The Overall Statistics application serves as the brief **summary of various statistical analyses** performed on the complete set of experimentally knwon phase separating protein sequences. The results are presented in the two-column layout, where on the left is the specified parameter and on the right is the corresponding total or average value. A broad overview includes many **chemical properties** (polarity, hydrophobicity, aromaticity, charge splited into positive and negative fraction), as well as other **biomolecular characteristics**, such as secondary structure (splited into helical and extended), solvent accessibility (splited into buried and exposed), structural disorder. All presented averages are derived from the consensus of the predictions of several state-of-the-art methods for a single LLPS sequence. Accordingly, the recently proposed **charge decoration parameters**, namely sequence charge decoration (SCD) and overall charge symmetry (OCS) together with a fraction of charged residues (FCR) are provided. There are also data indicating the actual number of BIAPSS-deposited LLPS-driver protein sequences, including the content of **disordered regions**, the number of organisms of origin and a **subset of human proteins**. Such a rough overview can give a glimpse into the LLPS protein set compared to other reference sets, while deeper analysis is further provided in the following interactive applications.

In separate tabs within the same web applications the user can find the **Length Distribution** and **Amino Acid Distribution**, and simultaneously compare the statistics performed for the complete LLPS sequences and extracted disordered regions.

*1)* The Length Distribution interactive analysis allows to simultaneously review the distribution of the length of LLPS sequences and the disordered fragments extracted from them in two switchable modes: presentation of lengths for individual sequences/disordered fragments (*distribution*) and *histogram* view. An interactive label in the distribution mode identifies the order index of the sequence/fragment in the database, the length of the sequence and its UniProt code, respectively. An interactive label in histogram mode identifies the range of the sequence length and counts of the corresponding sequences. The selected part of the chart can be easily enlarged with the automatic Plotly buttons on the top right.

*2)* The Amino Acid Distribution interactive analysis allows for a simultaneous comparison of fractions of each of the 20 biogenic amino acids in a set of LLPS sequences and structurally disordered LLPS fragments. There are 3 display modes to choose from: *distribution*, where an interactive label indicates the fraction of a given amino acid in a given LLPS sequence; *average*, where an interactive label indicates the average and standard deviation of the fraction of a given amino acid in the LLPS sequences set; *both*, where the average value is shown in relation to the distribution of fractions.

A comprehensive statistical analysis on a set of sequences prone to phase separation was carried out on 3 levels:

- entire LLPS sequences
- structurally disordered fragments of LLPS sequences
- short 20 amino acid segments along the LLPS sequences

Our approach to LLPS set provides a broad overview of the general biophysical characteristics and regularities which are common for known phase separating protein sequences. This makes it possible to infer certain overall regularities that may be helpful not only in efficiently identifying phase-separation-prone sequences but also for designing modifications for controlling LLPS.

#### *Amino Acid Composition App*

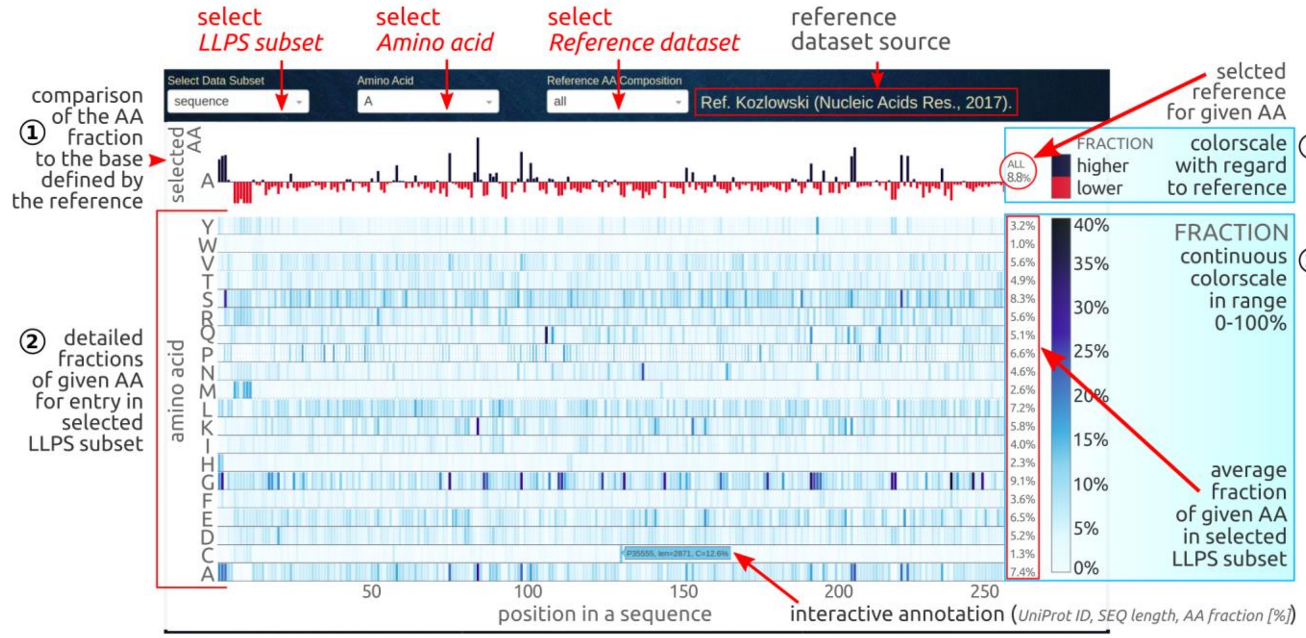


Figure S11. Snapshot of the AA Composition web application with its main functionalities highlighted by red arrows.

The Amino Acid Composition application enables the analysis of **amino acid composition** within the superset of LLPS-driver protein sequences. In the single interactive graph, the user can compare the **fractions of the 20 biogenic amino acids. The comparison is done** simultaneously for all LLPS-driver protein sequences deposited in BIAPSS repository. This additional customization facilitates comparison with the benchmark statistics derived for the other proteins grouped by organism of origin or protein type (globular, transmembrane, disordered).

##### *Dropdown options*

The user can select any of the following subsets of data: (i) the **full-length LLPS sequences**, (ii) short (<20 amino acids) and (iii) long (≥ 20 amino acids) LLPS **disorder fragments**. The second dropdown allows to select a particular AA type to compare its fractions with chosen benchmark statistics for other group of proteins or organism kingdoms (listed in the third dropdown).

##### *Graph section*

*1)* The top row is the content of the given amino acid type (2nd dropdown) for all entries from a selected data subset (1st dropdown) related to a benchmark fraction of same amino acid in reference dataset of proteins (3rd dropdown). The black and red bars indicate a **higher and lower fraction of the selected amino acid referred to selected dataset**. This supports the identification of trends in amino acid composition specific for phase-separating or disordered proteins. The reference content of the chosen amino acid for a selected reference set is given on the right as a percentage value and changes interactively when another amino acid type or reference set is selected.

*2)* The next multi-row section contains the **fractions of individual biogenic amino acids** in the selected LLPS data set, compiled at a single view. The fractions for the individual amino acids are aligned vertically per index (UniProt identifier) of LLPS sequence and displayed parallel in the multi-row layout. The attractive heatmap type of plotting allows for compact and straightforward all-in-one view by highlighting the cases exceedingly rich in a given type of amino acid relative to the complete set. The average fraction of a given AA in a selected data subset, is provided in the end for each row. A detailed annotation is available for the interactive labels.

#### *Chemical Properties App*

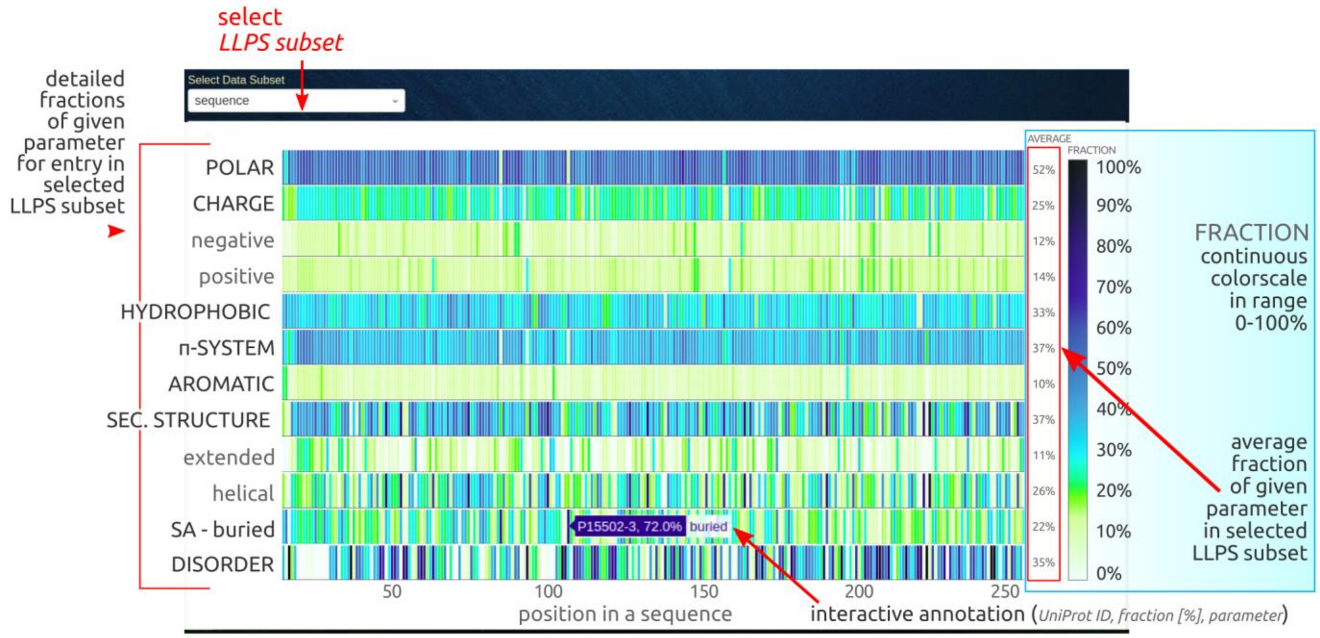


Figure S12. Snapshot of the Chemical Properties web application with its main functionalities highlighted by red arrows.

The Chemical Properties application summarizes the **chemical properties** and **biomolecular characteristics** on a single chart, for both the full-length LLPS sequences available in BIAPSS repository and for the LLPS disordered fragments. Specifically, the user can investigate the overall distribution of various chemical properties, such as **polarity**, **hydrophobicity**, **charge** (total, positive, negative), **aromaticity**, and **other π-systems**. Also, the condensed results of deeper sequence-based predictions are provided, including **secondary structure** (total, helical, extended), **solvent accessibility** (buried) and **structural disorder**. Compiling all the properties on a single chart, we make it possible to find certain correlations not only at the level of general trends for LLPS proteins, but also in a comparative analysis or a case study.

##### *Dropdown options*

The user can select any of the following subsets of data: (i) the **full-length LLPS sequences**, (ii) short (<20 amino acids) and (iii) long (≥ 20 amino acids) LLPS **disorder fragments**.

##### *Graph section*

The multi-row graph contains the chemical characteristics and sequence-based predictions for all of the entries from the selected LLPS data subset. For each sequence (or disordered fragment) indexed by UniProt ID, all of the individual parameters are calculated as the fraction of the residues in the sequence that meets the given criterion. Accordingly, the average value of each parameter across the entire LLPS subset is given at the end of each row. This can serve as the reference point for a particular case study. Further, compiling all of the properties for all of the sequences on the heatmap colored by the fractions, make it easier to find the correlation not only at the level of general trends for LLPS proteins, but also in a comparative analysis between cases. The interactive label indicates the UniProt ID for the selected sequence and percentage of the residues meeting a given property criteria.

#### *Charge Decoration App*

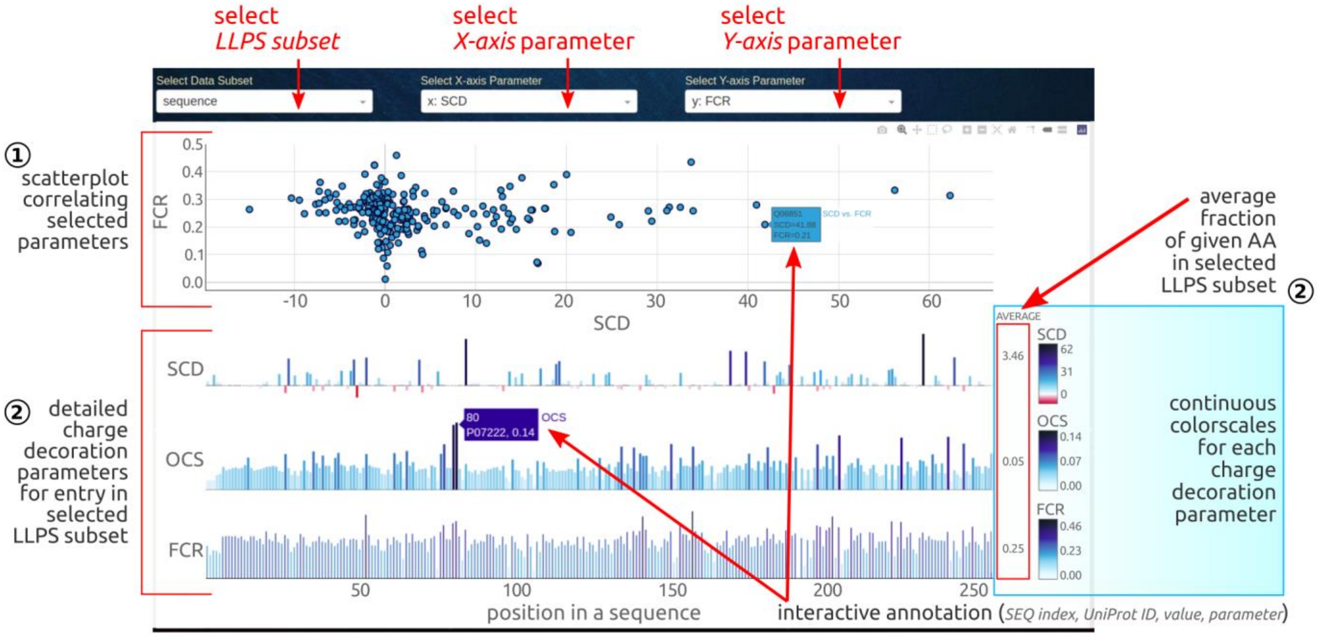


Figure S13. Snapshot of the Charge Decoration web application with its main functionalities highlighted by red arrows.

The Charge Decoration application is designed for an interactive comparison of various **charge decoration parameters** of the full-length LLPS sequences and the extracted LLPS disordered fragments. Specifically, the descriptors include sequence charge decoration (**SCD**), overall charge symmetry (**OCS**), and fraction of charged residues (**FCR**). The definitions of parameters are provided in the Docs tab of BIAPSS web platform. In the app the user can investigate the overall charge decoration patterns and follow the correlations on a complete set of LLPS proteins. The proposed parameters recently emerged as a good measure of charge distribution along the protein sequence, which, in addition to the overall charge content, turned out to be an important factor shaping the protein conformation, especially within low-complexity regions. Additionally, electrostatic interactions affect the solubility and stabilize the binding interface, and thus may appear to be relevant for the phase separation mechanism.

##### *Dropdown options*

The user can select any of the following subsets of data: (i) the **full-length LLPS sequences**, (ii) short (<20 amino acids) and (iii) long (≥ 20 amino acids) LLPS **disorder fragments**. The second and third dropdown menus allow correlating the two selected parameters on the scatter plot in the top section.

##### *Graph section*

*1)* In the top section it is possible to directly **compare two selected parameters** across the collective set of full-length sequences or extracted disordered regions.

*2)* The bottom section consists of three parallel bar charts showing **detailed distributions** of all descriptors. Since these measures are a single value per sequence, the background created by the distribution of values for a set of sequences becomes a good reference base in the analysis. The interactive label indicates the order index of the sequence in the database, UniProt ID for the selected entry and value of a given parameter.

#### Short Motifs – SET App

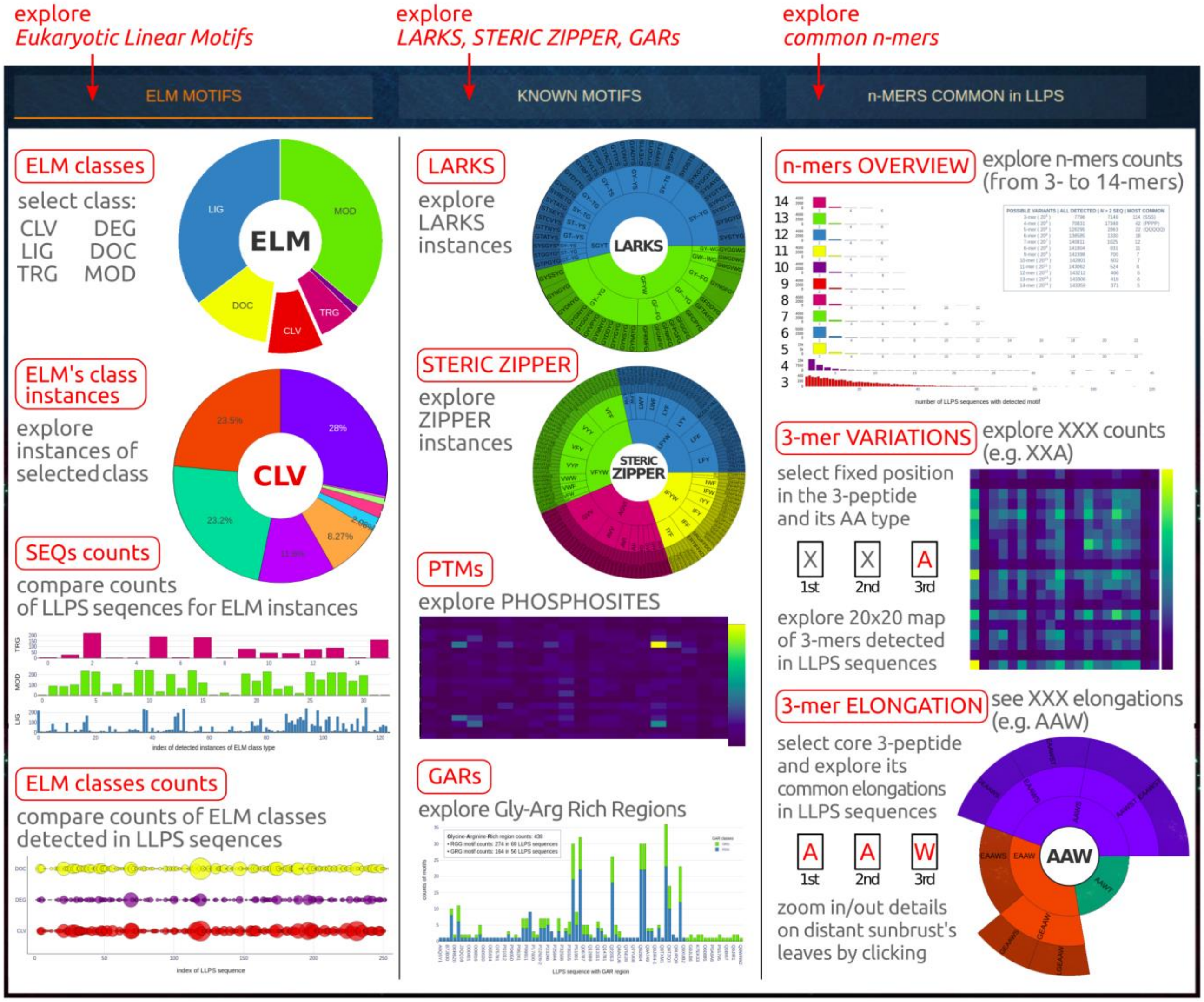


Figure S14. Snapshot of the Short Motifs - SET web application with its main functionalities highlighted by red arrows.

The application combines three main tabs providing the statistics on the detected SLiMs, including; (i) **eukaryotic linear motifs** (ELMs), (ii) other known short stretches of protein sequence such like **LARKS**, **Glycine-Arginine Rich regions** (GARs), **steric zippers** or **phosphosite motifs**, (iii) **novel motifs** filtered out as common from the systematic analysis of various n-mers. The implemented algorithms use the list of motif instances grouped and encoded by regular expressions as keys to search known phase-separating sequences. The interactive charts allow the user to choose between classes of motifs and show the total counts and occurrence in the individual LLPS sequence.

SLiMs have some unique properties and roles, such as evolutionary conservation, high structural flexibility, involvement in regulatory processes through the intermolecular interactions, that seems to be highly significant for phase separation. Encouraged by this lead, we carefully screened the LLPS sequences to detect not only the known SLiMs but also performed a systematic analysis of various 3 to 14-mers to carve out new ones.

#### *ELM MOTIFS tab*

**Eukaryotic Linear Motifs** are short stretches of adjacent amino acids experimentally validated in eukaryotic cells that are enriched in intrinsically disordered proteins and are crucial for compact intermolecular interactions. The list of functional sites was downloaded from the ELM server as a list of experimentally confirmed instances assigned to each of the 6 main classes:

- CLV - cleavage sites,
- DEG - degradation sites,
- DOC - docking sites,
- LIG - ligand binding sites,
- MOD - post-translational modification sites,
- TRG - targeting sites.

The user can select between pie, bar, or bubble charts to display ELM data.

*1) Pie chart options.* The app-page consists of two interactive pie charts:

- on the left - the graph shows the percentage share of individual classes (CLV, DEG, DOC, LIG, MOD, TRG) in the total number of ELMs detected in LLPS sequences.
- on the right - the graph shows the percentage share of individual instances in the total number of selected ELM class detected in LLPS sequences.

The interactive labels indicate the ELM motif class/instance and its counts. An interactive legend permits excluding individual classes or instances from the statistics.

*2) Bar chart options.* The app-page consists of a 6-row bar chart, where the single row is dedicated to instances of selected ELM class. The buttons allow determining the total frequency of specific classes of ELM motifs, including detailed counts of individual instances and the number of unique LLPS sequences in which the motifs were detected.

*3) Bubble chart options.* The app-page consists of a 6-row bubble chart, where the single row is dedicated to instances of selected ELM class. The bubble's size determines the total counts of specific classes of ELM motifs detected in each unique LLPS sequence. The comparative analysis supports the identification of ELM's richest sequences as well as the detection of the co-existence of various motifs.

#### *KNOWN MOTIFS tab*

Recent studies indicate many other short motifs along the sequence, which have tendency to engage in multivalent interactions broadly suggestive of the involvement in phase separation or aggregation:

- **LARKS** are the low-complexity, aromatic-rich, kinked segments, where the experimental studies showed that the two of such short sequence fragments could bind weakly to each other by forming a pair of kinked β-sheets.
- **steric zipper** is a structural motif of the dry interface formed by the interdigitation of complementary side chains via van der Waals interactions and hydrogen bonds such as Asn/Gln ladders.
- **GARs** are Glycine-Arginine-rich disordered protein regions with multiple RGG/RG repeats that may bind nucleic acids via weak multivalent interactions.
- **PTM** is a modification of proteins that increases the functional diversity of the proteome by the covalent addition of various functional groups.

The user can explore various charts to display KNOWN motifs data.
*1) Pie chart options.* The pie chart for each of LARKS, steric zippers, and phosphosites shows the counts of instances for the generalized pattern of motifs and a list of UniProt IDs of LLPS sequences for which the particular instances were detected. The interactive on-click events allow for customized zooming in and out of the user-selected instances.

*2) Heatmap chart options.* The heatmap charts for phosphosite motifs allow users to track the frequency of specific instances of the motif according to their amino acid composition at the second and third positions in the sequence fragment.

*3) Stacked Bar chart options.* The stacked bar chart for Glycine-Arginine Rich regions compares the distribution and counts of the two common instances of GARs (GRG/RGG) in the LLPS sequences.

#### *n-MERS tab*

Known Linear Motifs are short fragments along the sequence that are generally evolutionary conserved and often situated in intrinsically disordered regions. Their unique feature is structural flexibility since the secondary structure is usually induced when interacting with a structured partner. This section contains results of our careful screening the LLPS sequences to detect various n-mers to help identifying novel motifs.

The user can explore:
*1)* **the overall distribution of n-mers** (from 3-mers to 14-mers) that are common across LLPS sequences.
The Bar chart shows the distribution of various n-mers which were found in at least 2 LLPS sequences. From the app, the user can learn about the most common short fragments of various lengths and explore detected instances. In the upper right corner, there is a brief summary of statistics, where the following parameters are provided:
• *possible variants* - total number of all possible fragments of length n created from 20 standard amino acids,
• *all detected* - number of unique fragments of length n that have been detected in at least one LLPS sequence,
• *n > 2 SEQ* - number of unique fragments of length n that have been detected in at least three LLPS sequences,
• *most common* - number of LLPS sequences in which the most common instance of n-mer was detected, e.g., SSS as the instance of 3-mers was detected in 144 LLPS sequences. The interactive on-hover annotations allow for a detailed analysis of the user-selected n-mers and their most common instances detected in the set of LLPS protein sequences.

*2)* **the core 3-mers and their elongated instances** (up to 6-mers) that can prove to be new short multivalent stretches of LLPS sequences.
The heatmap chart allows users to track the frequency of specific instances of the 3-mer according to their amino acid composition. Using the available options, the user can select the fixed position in the 3-peptide (XXX) and its amino acid type, while the amino acids at two remaining positions are encoded on a 2-dimensional map. In-depth analysis of the user-selected 3-peptide and their most common elongation up to 6-mers may bring the pioneering discovery of novel short linear stretches.
When an interesting 3-mer (e.g., very common in LLPS sequences) is detected, the user can use it as an input to investigate more deeply the detected elongations of a core 3-peptide in a set of LLPS sequences. This is possible on the right in the sunburst chart. The interactive on-hover labels include core 3-peptide type, elongated motif instance, number of LLPS sequences with detected motif and list of UniProt IDs, total counts.

#### Local Fragments App

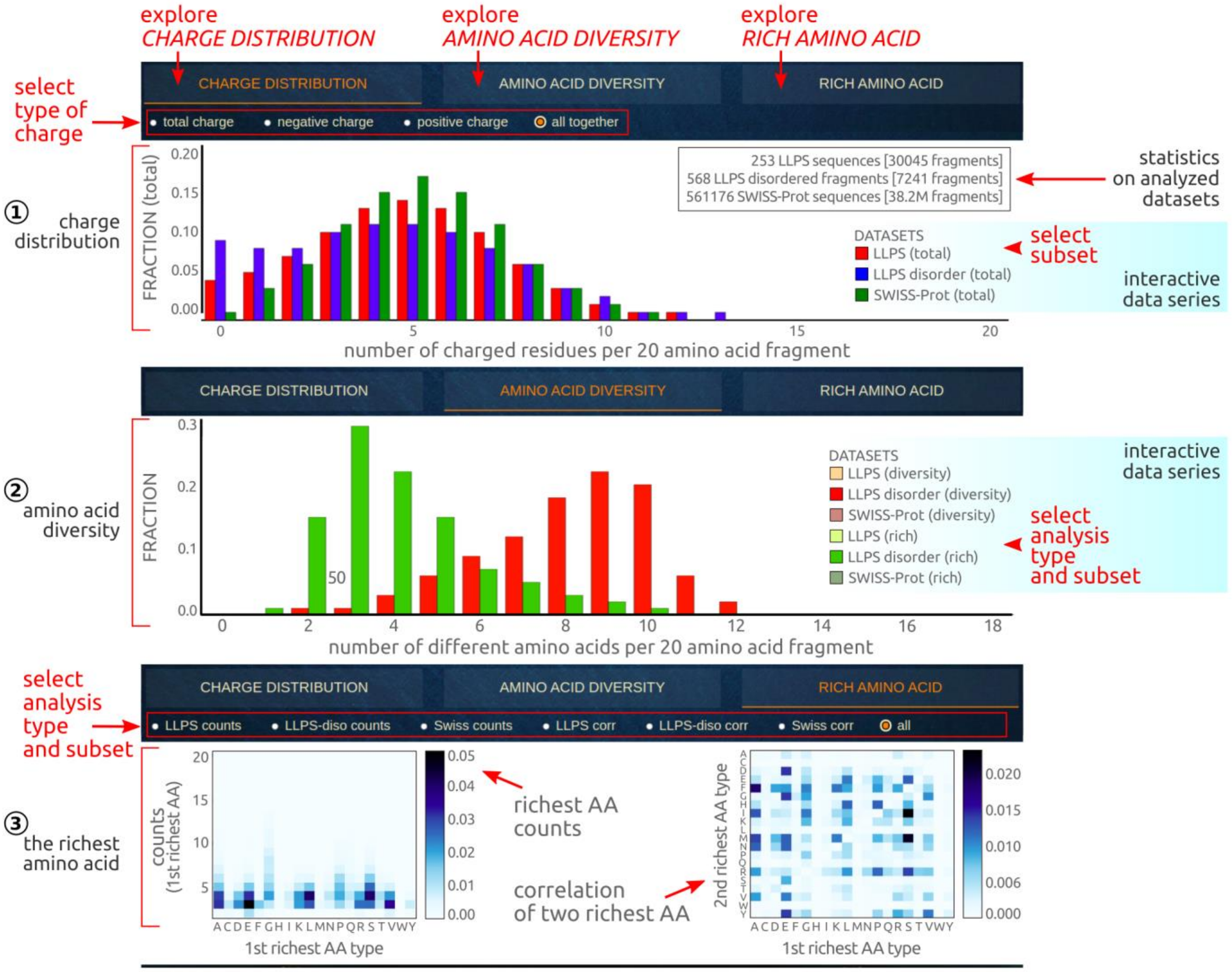


Figure S15. Snapshot of the Local Fragments web application with its main functionalities highlighted by red arrows.

A rough statistical analysis of the full-length sequences, described by a set of averages or fractions, gives general insight but conceal some local peculiarities and make certain attributes utterly invisible. For this reason, we conducted additional analysis on short 20 amino acid long fragments extracted along the LLPS sequences with the 5 residues shift. The central aim of the study was to identify some local regularities relevant for phase separating proteins as well as learn the pattern of their organization.

The Local Fragments application combines the results of comprehensive analysis conducted on the **20-residue fragments** extracted from the full-length LLPS sequences (253), structurally disordered LLPS regions (568), and sequences from the SwissProt set (561176, used as a reference benchmark). For obtained fragments we examined (i) the **distribution of charges** (total, positive, negative), (ii) and **amino acid compositional diversity**, (iii) also we indicated **AA-rich local repeats**.

The web application contains the results of 3 comprehensive analyses:

*1) CHARGE DISTRIBUTION*, contains the charge distribution. The user can select the displayed data using any subset inclduing features such as: total charge, positive charge, negative charge, or all together. An interactive legend allows switching between the displayed LLPS data subset. The interactive label indicates the number of charged residues per 20 amino acid fragment and the fraction of residues with a particular type of charge. The top right text area describes the counts of analyzed sequences and 20 AA fragments.

*2) AMINO ACID DIVERSITY*, compares the AA compositional diversity which is measured by two parameters:

• *AA diversity* - the number of diverse residues observed within 20 amino acid fragments,

• *AA uniformity* - the counts of the common amino acid residue observed within 20 amino acid fragments.
An interactive legend allows to switch between the displayed data subsets. The interactive label indicates the number of eligible residues per 20 amino acid fragment and the corresponding fraction (counts) of fragments meeting the condition.

*3) RICH AMINO ACID*, contains the detailed analysis of the richest amino acid residues observed within 20 amino acid fragments from all of the benchmark datasets. Within this tab the user can detect the residues which are enriched in the LLPS sequence set as well as learn the pattern of these residues (counts per 20 AA fragment). The user can select data subsets and display results in terms of the *counts* and *correlation* modes, or see all of the charts together (default mode). The interactive label contains detailed scores. The heatmap charts correlate two the most abundant amino acids in the 20-residue long fragments (counts of each higher than 1) and indicate the pair of amino acids prone to be closely located in the sequence. These findings can help to identify interactions relevant to the phase separation.

#### Download TAB – Repository of pre-computed data and statistics

availability: *https://biapss.chem.iastate.edu/download.html*

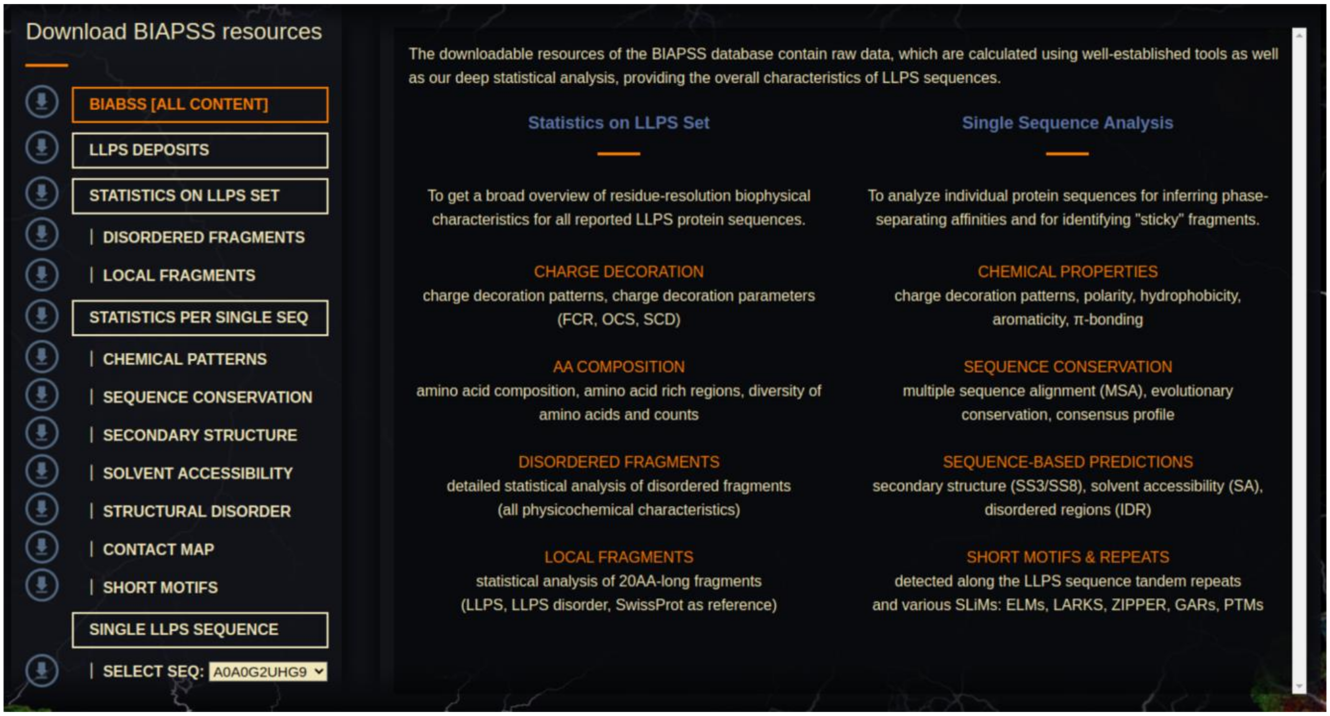

Figure S16. Snapshot of the Download section of the BIAPSS web platform.

Download tab makes the BIAPSS pre-computed repository freely accessible for users. Its main aim is to store compressed and ordered data files by using the straightforward CSV format. These pre-processed files are designed to reduce the complexity of BIAPSS entries as well as to provide fast access to results of high-quality sequence-based analysis coupled with additional features such as rich annotations to external databases and agreed consensus of predictive methods. The available data includes raw predictions pre-calculated using well-established tools as well as findings of our deep statistical analysis, giving the overall characteristics of LLPS sequences. For the user convenience, we unified and integrated the results from these sources into several categories by following the list of applications in *SingleSEQ* and *MultiSEQ* tabs.
